# Keystone species determine the productivity of synthetic microbial biofilm communities

**DOI:** 10.1101/2022.01.23.477386

**Authors:** Xinli Sun, Jiyu Xie, Daoyue Zheng, Riyan Xia, Wei Wang, Weibing Xun, Qiwei Huang, Ruifu Zhang, Ákos T. Kovács, Zhihui Xu, Qirong Shen

**Author notes:** Corresponding authors: Zhihui Xu and Ákos T. Kovács. Xinli Sun and Jiyu Xie contributed equally to this work.

## Abstract

Microbes typically reside in multi-species communities, whose interactions have considerable impacts on the robustness and functionality of such communities. To manage microbial communities, it is essential to understand the factors driving their assemblage and maintenance. Even though the community composition could be easily assessed, interspecies interactions during community establishment remain poorly understood. Here, we combined co-occurrence network analysis with quantitative PCR to examine the importance of each species within synthetic communities (SynComs) of pellicle biofilms. Genome-scale metabolic models and *in vitro* experiments indicated that the biomass of SynComs was primarily affected by keystone species that are acting either as metabolic facilitators or as competitors. Our study sets an example of how to construct a model SynCom and investigate interspecies interactions.

## Introduction

Despite a wealth of information available on the microbiota composition gathered from sequencing techniques, little is known about the interspecies interactions that govern their assemblage and dynamics. Yet, the size and complexity of natural microbial communities are often too large to be manipulated. Synthetic microbial communities (SynComs) have been proposed as model systems to overcome these challenges (Großkopf and Soyer, 2014). Several SynComs with moderate complexity and high controllability have been developed to represent different natural environments, such as plant rhizosphere (Bai et al., 2015; Ma et al., 2021; Niu et al., 2017; Schmitz et al., 2022; Voges et al., 2019), phyllosphere (Berg and Koskella, 2018; Carlström et al., 2019), and intestine (Hromada et al., 2021; Ortiz et al., 2021; Weiss et al., 2021). In nature, most bacteria aggregate as sessile multicellular communities that are embedded in the self-produced extracellular matrix called biofilm (Flemming and Wuertz, 2019; Webb et al., 2003). Biofilm communities have emergent properties that are not predictable from free-living bacterial cells (Flemming et al., 2016), including enhanced biomass production (Ren et al., 2015), stress tolerance (Lee et al., 2014; Orazi and O’toole, 2019), and improved resource acquisition (Nielsen et al., 2000). The spatial organization of biofilms facilitates the intermixing of cooperating species and spatial segregation of competing species (Nadell et al., 2016). Understanding how bacterial interactions affect the assemblage of bacterial communities has direct applications in biotechnology, agriculture, and health (Bengtsson-Palme, 2020; Cavaliere et al., 2017; Cho and Blaser, 2012; Fitzpatrick et al., 2020; Gómez-Godínez et al., 2021).

Studies on SynComs reported that the assemblage and robustness of communities can be affected by several factors including pH (Ortiz et al., 2021), temperature (Burman and Bengtsson-Palme, 2021), spatial distribution (Liu et al., 2019), initial abundance (Gao et al., 2021), niche specificity (Estrela et al., 2021), nutrient availability (Ratzke et al., 2020), and keystone species (Niu et al., 2017). Five principles need to be considered when developing SynComs: representativity, stability, reduced size, accessibility, and tractability (Blasche et al., 2017). To assemble a representative and stable SynCom, microbial co-occurrence network analysis could serve as guidance for the selection of isolates (Poudel et al., 2016). Positive or negative associations indicate candidate taxa. This method has been applied to investigate factors affecting host microbiome variation (Agler et al., 2016), to link taxa to biological functions of interest (Wei et al., 2019), to identify potential biotic interactions (Durán et al., 2018), and to explore habitat differentiation (Barberán et al., 2012). However, the ecological relevance of predicted interactions remains poorly understood (Faust, 2021). The connectedness and strength of positive or negative interactions are not experimentally verified. Another emerging method for exploring microbial interactions is genome-scale metabolic modeling, which can provide insights into metabolic interaction potential and metabolic resource overlap in multi-species communities (Zelezniak et al., 2015a; Zorrilla et al., 2021). Accessibility and tractability require a community to contain limited members, and the community members are cultivatable and can be promptly and accurately quantified. This can be achieved by colony-forming unit (CFU) counting (Niu et al., 2017; Piccardi et al., 2019), fluorescent labeling (Kehe et al., 2021), and quantitative PCR (Ren et al., 2015). While CFU counting is simple to employ but time-consuming, the latter two approaches are more efficient but require extensive initial preparation time and effort. Reduced size permits the manipulability of a SynCom.

In this study, we constructed multi-species biofilm communities using isolates from a rhizosphere and predicted their interactions by analyzing co-occurrence networks. We evaluated the role each member played in community productivity and its impact on the other members of the community. The potential metabolic interactions were further investigated through experiments and metabolic modeling. Our results suggested that keystone species determine community productivity, probably through metabolic exchanges or resource competition. We propose that our study could inform the rational design of synthetic communities.

## Results

### Source of SynCom members

As co-existence facilitates biofilm formation in multi-species communities (Madsen et al., 2016), we adopted co-existence as the first criterion of SynCom selection. To assemble a soil biofilm community, we co-cultured the rhizosphere soil with *Bacillus velezensis* SQR9 (a strong biofilm former widely studied in our lab, abbreviated as Bac) to form pellicle biofilms (Figure 1—figure supplement 1A). We determined the bacterial composition of the biofilm and the solution underneath using the amplicon sequencing method. Consequently, 15 genera and 2 families were predicted to co-exist in the culture condition (Figure 1—figure supplement 1B). From our bacterial collection (Sun et al., 2021), we selected 11 isolates that corresponded with the predicted co-existing taxa to represent the soil biofilm community (Figure 1—figure supplement 2). These isolates are derived from three different phyla: Firmicutes, Proteobacteria, and Bacteroidetes.

Generally, biofilm development undergoes various stages including initial motility, development, maturation, and disassembly (Vlamakis et al., 2013). To evaluate the stability of the SynCom, we expanded the volume of cultivation to 400 ml and tracked the dynamic changes of bacterial composition: development (2d), early maturation (4d), late maturation (6d), and disassembly (8d). The bacterial composition stabilized at the maturation stages. Chr (*Chryseobacterium rhizoplane*) was identified to be the most predominant member followed by Aci (*Acinetobacter baumannii*) as the second most abundant species throughout the biofilm formation process (Figure 1A). Three isolates Ach (*Achromobacter denitrificans*), Pxa (*Pseudoxanthomonas japonensis*), and Ste (*Stenotrophomonas maltophilia*) rapidly declined and could not establish themselves during biofilm maturation (4d & 6d). At the biofilm disassembly stage (8d), the sticky, robust biofilm structure has dispersed as small, fragile aggregates. Most cells have lysed, thus the relative abundance of rare isolates increased.

**Figure 1.**
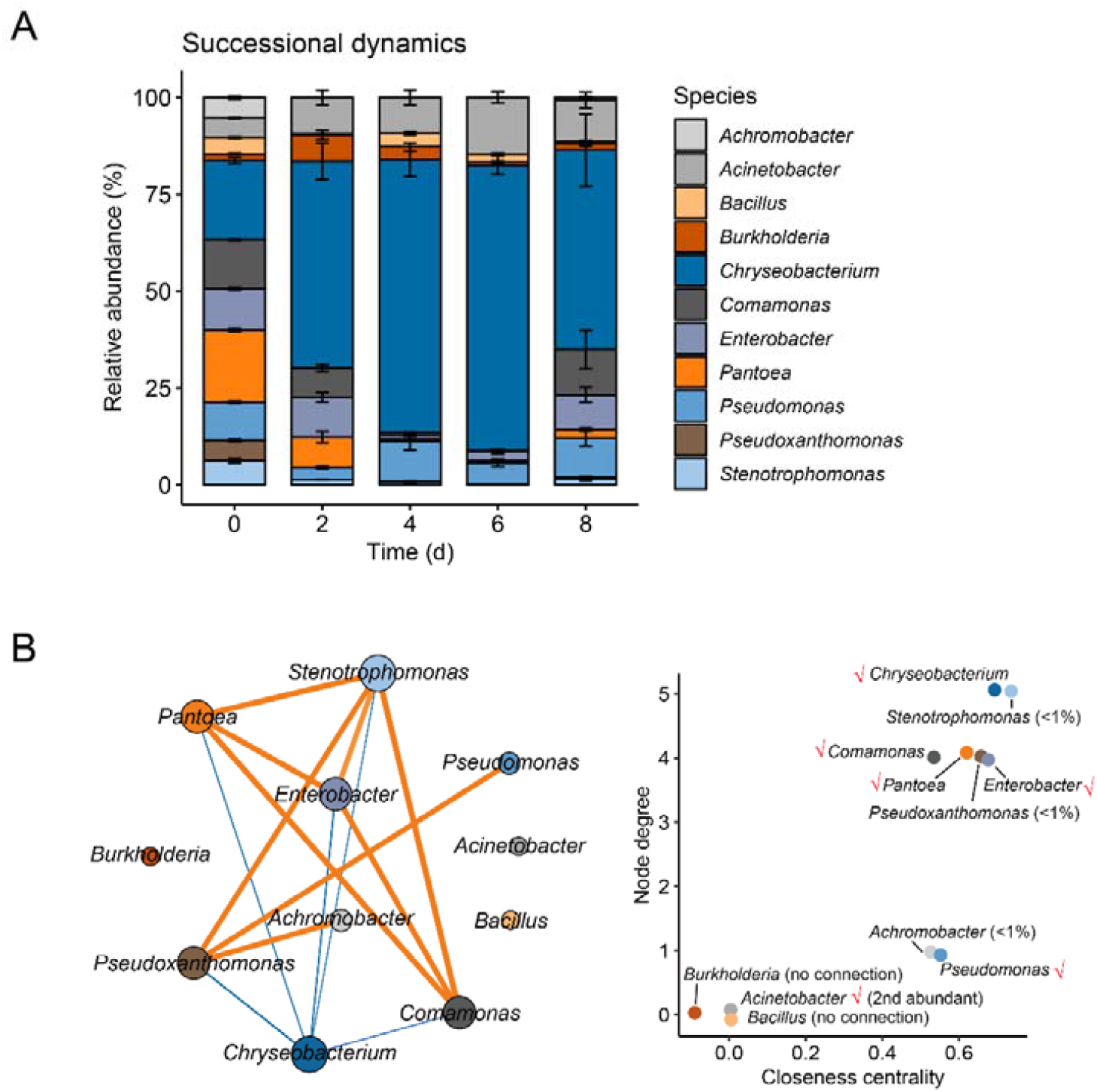
Dynamic changes of the initial 11-species consortium composition in the biofilm. **(A)** The relative abundance of isolates determined by 16S rRNA gene amplicon sequencing method. Data presented are the mean ±sd. n =8. **(B)** Co-occurrence network of bacteria within biofilm communities. Each node represents a bacterial species, node size is proportional to node degree. Line width indicates the interactive strength of interaction, and color indicates the sign of correlation (orange indicates positive, blue indicates negative). A positive correlation means that as one isolate increase, the other isolate also tends to increase, and vice versa. **(C)** Selection of six isolates. Node degree is the number of direct correlations to a node in the network. Closeness centrality indicates the average distance from a given starting node to all other nodes in the network. Ticks indicate isolates selected in the following reduced community. These isolates were selected based on either high abundance or high correlations as indicated by node degree and closeness centrality. **Figure supplement 1**. Origin of the initial 11 isolates. **Figure supplement 2**. A flow chart of isolates used for the study, their origin and selection criteria.

We hypothesized that microbial interactions could contribute to the compositional changes. Therefore, network co-occurrence analysis was employed to infer the correlations (Spearman’s correlation coefficient *r* > 0.6, *p* <0.01) among the species during biofilm formation (Figure 1B). Co-occurrence network revealed 9/14 positive correlations and 5/14 negative correlations. Intriguingly, the negative correlations all involve Chr, suggesting that the increment of Chr is linearly correlated with the reduction of other isolates. Aci, Bac, and Bur (*Burkholderia contaminans*) had no connection with others, indicating that they had little influence on the growth of other isolates (Figure 1B). The connectedness of isolates was further evaluated by node degree and closeness centrality (Figure 1C). Node degree is the number of edges the node has. The higher the degree, the more central the node is. High closeness centrality indicates the node is closely connected to other nodes and is central in the network. As a result, six isolates - Chr, Ste, Com (*Comamonas odontotermitis*), Pan, Ent (*Enterobacter bugandensis*), Pxa - are closely connected with other isolates and are central in the network. Ach and Pse are closely connected with other nodes but not central in the network.

In the subsequent experiments, we selected six isolates to examine bacterial interaction. They were selected based on high relative abundance (Chr, Aci) or high closeness centrality (Pan, Com, Ent, Pxa, Pse). Chr fulfilled both criteria and was hypothesized as the keystone negative species in the reduced SynCom. Other species were excluded either due to their extremely low abundance at the biofilm maturation stage (Ach, Pxa, Ste), or due to the lack of correlations with the other species (Bac, Bur).

### The productivity of multi-species biofilm was affected by central nodes

The co-occurrence network suggested Pan, Com, and Ent are positively correlated with each other, while they are all negatively correlated with Chr. Aci and Pse are not connected with the selected nodes. We hypothesized that the removal of central nodes (Pan, Com, Ent, Chr) would have a significant impact on community structure and productivity, while the removal of non-connected nodes (Aci, Pse) would have no impact. To evaluate the importance of each isolate on biofilm productivity, we applied the “Removal” strategy: the “Full” community consists of six isolates, then one isolate was dropped out to obtain the five-species reduced communities, abbreviated as “Rm” communities. Biofilm productivity was assessed in two ways: fresh weight and population cell numbers (Figure 2A). We minimized the volume of biofilm cultivation to 10 ml to allow massive parallel sampling of the different combinations. Importantly, the smaller volume shortened the biofilm development time, thus two earlier time points were tested: 24h represents the biofilm development stage, while 36h represents the maturation stage. The cell numbers were quantified by strain-specific primers (Table 1). The role of each isolate is represented as changes in community productivity by removing the certain isolate from the full community.

**Fig 2:**
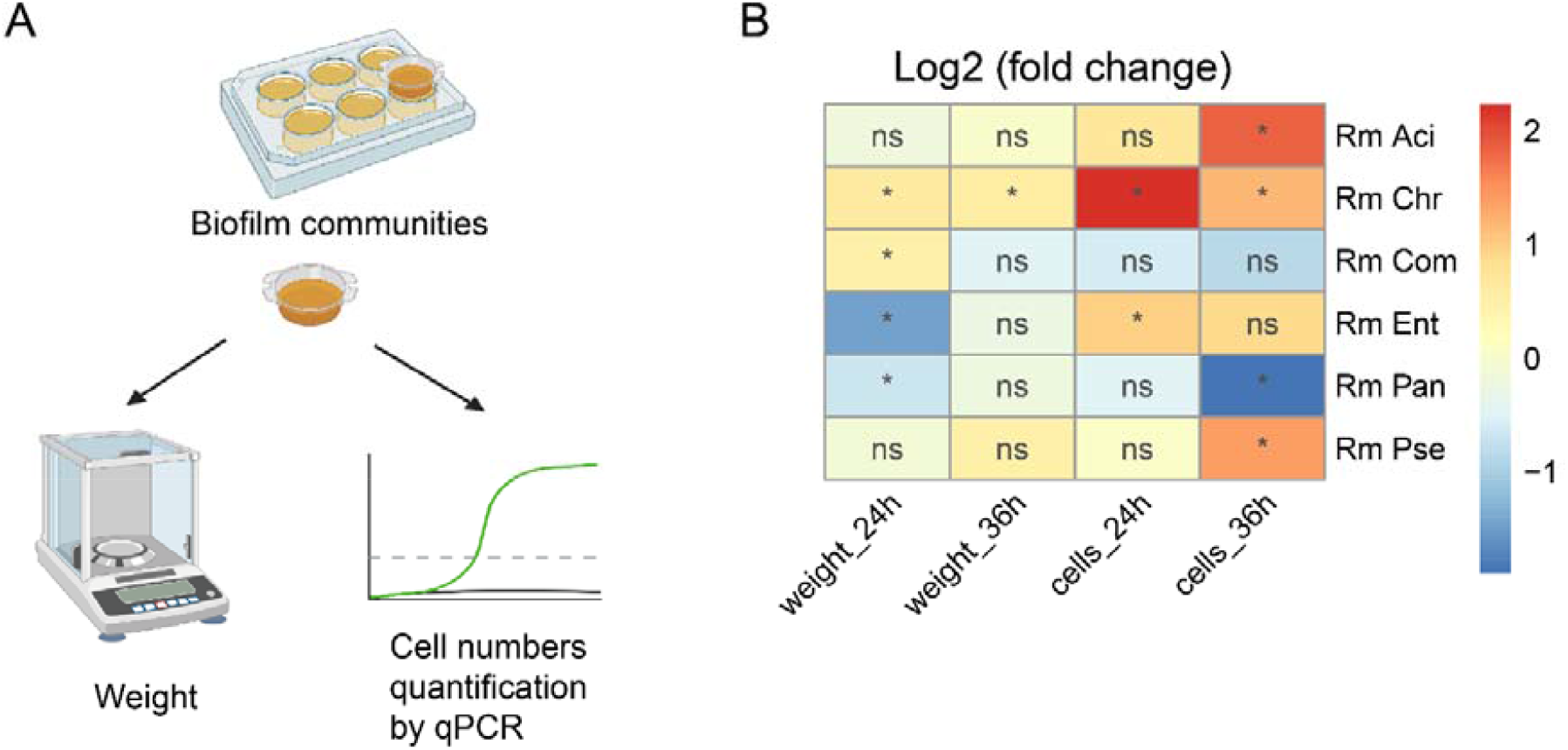
Productivity of the reduced SynComs. **(A)** Experimental design. **(B)** Changes in biofilm weight and population cell numbers. “*”indicates the value of the reduced community is significantly changed compared with the “Full” community (ANOVA, Tukey’s test, *p* <0.05). Values of the replicates are shown in figure supplement 1.Rm Aci, Rm Chr, Rm Com, Rm Ent, Rm Pan, Rm Pse represent the five-species communities resulting from the removal of *Acinetobacter baumannii* XL380, *Chryseobacterium rhizoplanae* XL97, *Comamonas odontotermitis* WLL, *Enterobacter bugandensis* XL95, *Pantoea eucrina* XL123 and *Pseudomonas stutzeri* XL272, respectively. **Figure 2 – figure supplement1**. Productivity of the reduced SynComs.

**Table 1.**
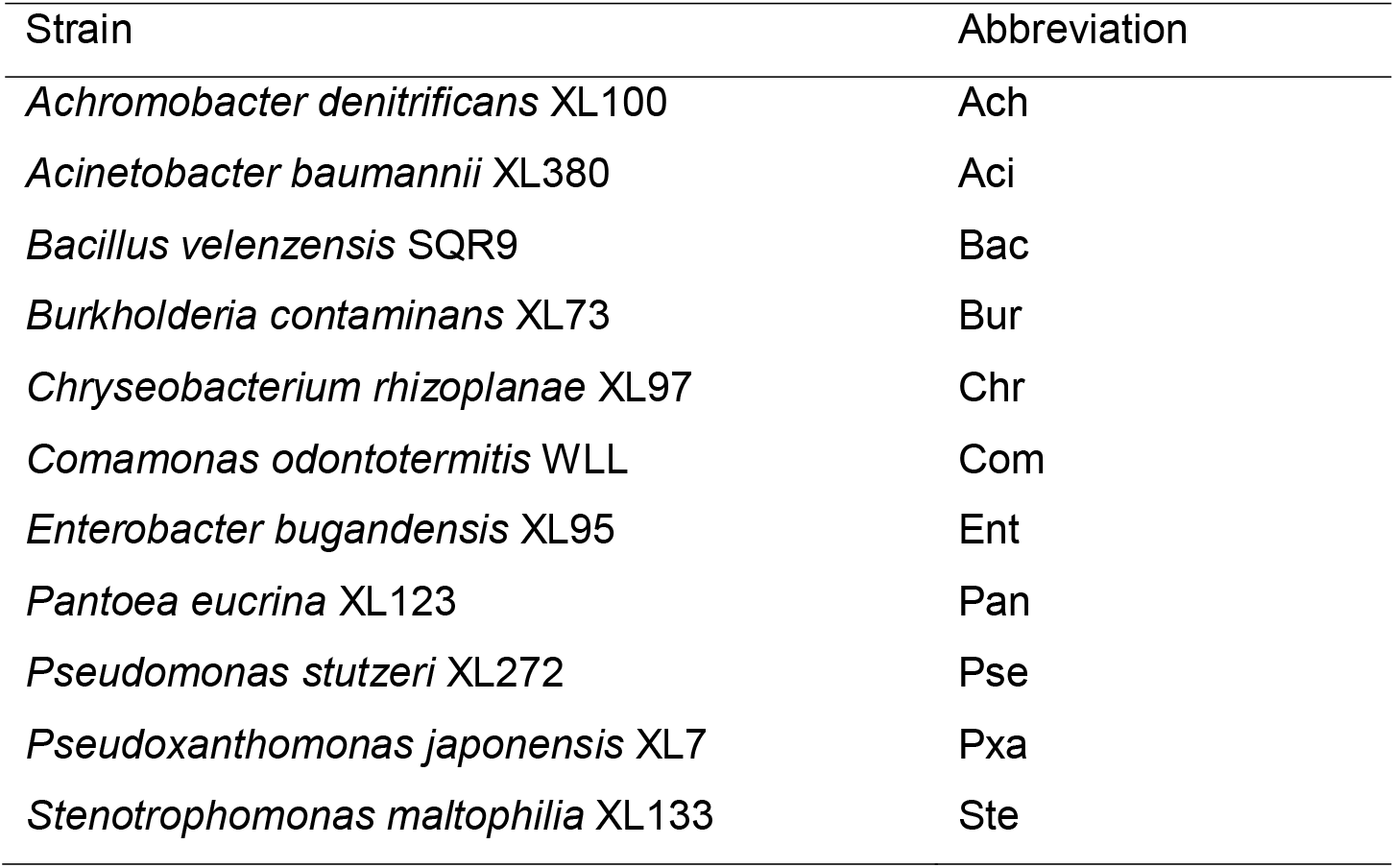
Strains used in this study

“Rm Chr” resulted in higher biofilm weight and higher community cell numbers at both time points (ANOVA, Tukey’s test, *p* <0.05) (Figure 2B & Figure 2 – figure supplement1). Conversely, “Rm Pan” led to lower biofilm weight at 24h and lower community cell numbers at 36h. Removal of the other two positive nodes (Com and Ent) did not result in the same effect. These partly confirmed our predictions: the negative node Chr and the positive node Pan affect the carrying capacity of SynCom. Besides, the removal of non-connected nodes (Aci and Pse) had little impact on community productivity, which also supports our hypothesis. The contradictory effect of “Rm Ent” on weight and cell numbers at 24h made it difficult to assign a clear role.

We also compared the individual cell numbers in the reduced SynComs with that in the “Full” community to evaluate the impact of removing one isolate on other isolates (Figure 3& Figure 3 – figure supplement). In general, most community members increased after the removal of Aci, Chr, or Pse, suggesting their presence of them inhibited the growth of other isolates in the SynCom. “Rm Pan” resulted in the reduction of Chr, Com, and Pse at 36h, suggesting Pan was important for supporting their growth at the biofilm maturation stage. “Rm Ent” also increased the cell numbers of Pan, indicating Pan was negatively correlated with Ent, which contradicted with co-occurrence network.

**Fig 3:**
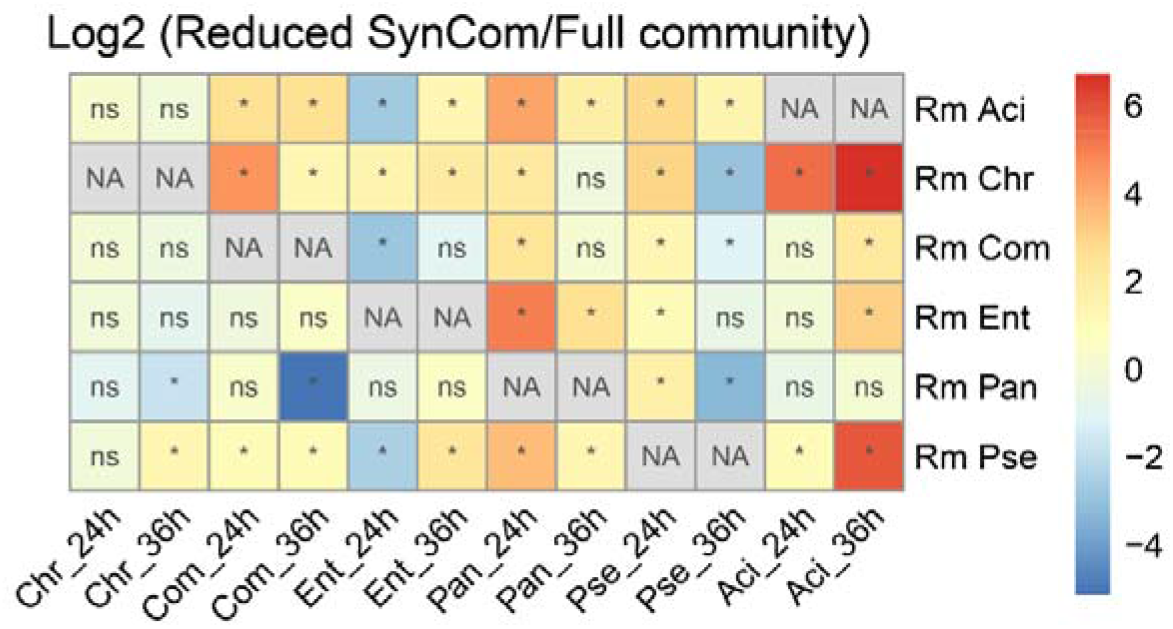
Influence of removing one isolate on SynCom composition. “*” indicates the cell numbers of the isolate in the reduced community is significantly changed compared with that in the “Full” community (ANOVA, Tukey’s test, *p* <0.05). Values of the replicates are shown in figure supplement. **Figure 3–figure supplement**. Individual cell numbers in different SynComs.

Taken together, we proposed Chr as a keystone negative species in the SynCom in terms of limiting the community productivity and inhibiting the growth of every other isolate in the community context. Pan is predicted to be a positive keystone species because its removal decreased the growth of three members and decreased the overall community productivity.

### Metabolic facilitation could explain the positive interaction

To gain insights into the metabolic interaction potential of the SynCom members, genome-scale metabolic models were established using the CarveMe pipeline. The M9 glucose minimal medium was set as the input medium because this medium can be experimentally validated and due to lack of available metabolic modeling pipeline for complex growth media, such as TSB. We derived the likely exchanged metabolites across communities and the strength of metabolic coupling - SMETANA score (Zelezniak et al., 2015). In the “Full” community, Pan served as the principal metabolic donor, all the other isolates received benefits from it (Figure 4A). On the contrary, Chr acted as the highest metabolic receiver, while Pan had the least potential to be the metabolic receiver. Amino acids, organoheterocyclic compounds, and phosphate were the major exchanged metabolites (Figure 4B). Aci and Com mainly receive amino acids from Pan, Chr, and Ent receive organoheterocyclic compounds from Pan, while Pse receives organoheterocyclic carbohydrates from Pan (Figure 4C).

**Fig 4:**
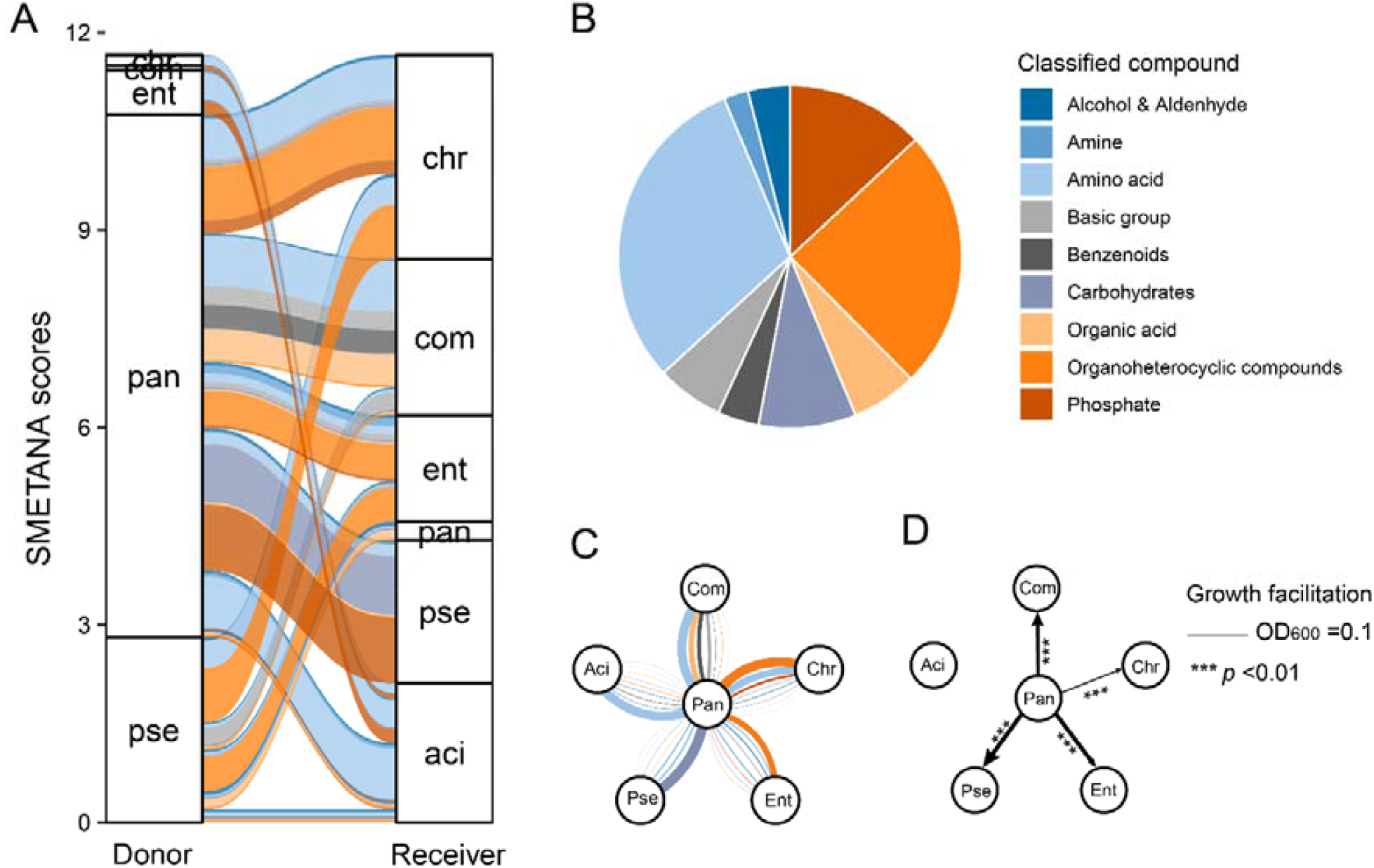
Metabolic facilitation. **(A)** Alluvial diagram showing the metabolic interaction potential of the “Full” community simulated by genome-scale metabolic modelling. Thickness of strips are SMETANA scores. **(B)** Pie chart showing the proportion of compounds exchanged within communities. **(C)** Flower plot showing metabolic interactions involving *P. eucrina* XL123 (centered in each panel) as a donor. Thickness of lines are proportional to magnitude of SMETANA score. Colors of all plots are the classification of compounds. **(D)** Growth facilitation assessed by growth in spent medium. The spent medium was prepared by growing the corresponding species in M9 medium with 0.2% glucose till the glucose was under detection then filter sterilized. The species were grown in the filtered spent medium for 24 h and the growth were measured as the OD_600_. Line width indicates the growth facilitation, that is the final OD_600_ substact the initial OD_600_. Only growth facilitation larger than 0.1 were shown in the figure. The values were compared with that of no inoculation control. \*\*\**p* <0.01, t test. **Figure 4 – figure supplement** Growth curves in the M9 spent medium

Growth assays were used to determine the accuracy of the metabolic models. Firstly, only four isolates were able to grow individually in the M9 medium with 0.2% glucose as sole carbon: Ent and Pse displayed high growing capacity, Aci and Pan grew moderately, while Chr and Com showed no growth (Figure 4 – figure supplement). Therefore, Ent, Pse, Aci, and Pan could have the potential to serve as metabolic donors in the M9 glucose medium. Secondly, growth in the spent culture medium was utilized to determine metabolic interaction potential. Each isolate was cultured in the M9 glucose medium until glucose was undetectable. The obtained sterile spent medium was used to cultivate every other member of the community and themselves. The maximum growing capacity was determined and defined as growth facilitation (Figure 4D). The spent medium of Pan was capable of supporting the growth of Aci, Chr, Ent, and Pse (Figure 4 & Figure 4 – figure supplement). These results might explain the positive role of Pan in the biofilm community. On the contrary, no isolates could grow in the metabolic by-products of Aci, Ent, or Pse. Chr could be a strong competitor for the metabolic by-products from Pan in SynCom.

### Resource competition could explain the negative interaction

Interspecies interactions in the SynComs may further be explained by metabolic competition. Using metabolic modeling, the metabolic resource overlap was simulated (Figure 5A). We observed high metabolic resource overlap across all communities, indicating intense resource competition among SynCom members. The removal of Chr minimized the metabolic resource overlap the removal of Pan or Pse resulted in higher resource competition. To test the possibility of direct competition, we performed spot-on-lawn and pair-wise spot assays on TSB agar plates (Figure 5B-C). Both Aci and Ent completely inhibited the growth of Com. Similarly, Chr and Ent inhibited Pse growth. However, these inhibitions were not caused by direct antagonism as no inhibition zones were observed in the respective spot-on-lawn assay plates (Figure 5B, red square). In addition, no clear boundary was observed between the two strains when they were spotted adjacent to one another (Figure 5C). Interestingly, Pan and Ent could change the colony color of Chr from light yellow to white (Figure 5B, yellow square).

**Fig 5:**
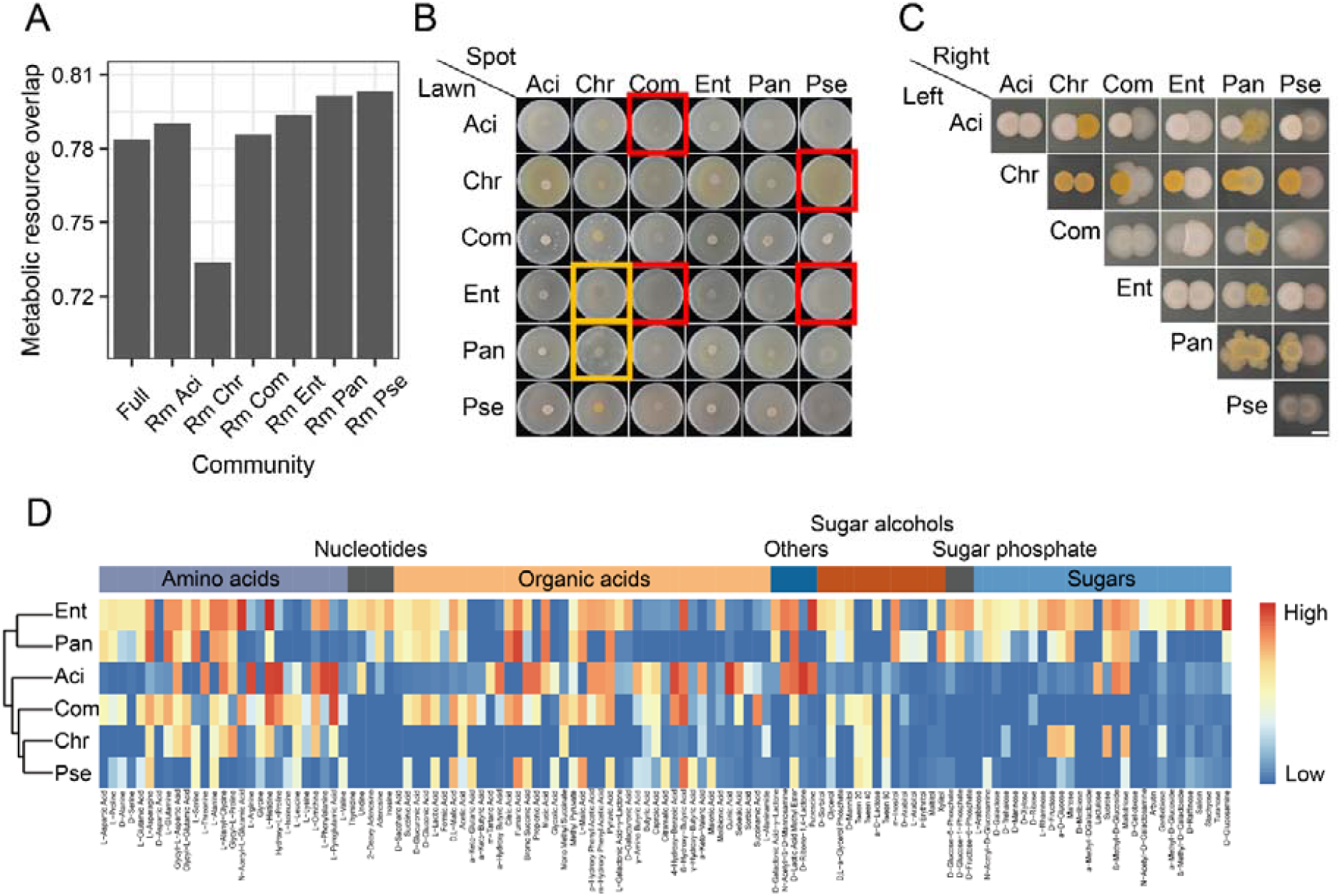
Resource competition. **(A**) Bar chart showing the metabolic resource overlap simulated by the metabolic modeling. **(B)** Spot-on-lawn assays. The lawn species were spread on TSB agar plates at an OD_600_ of 0.02 and dried, 5 μl of the spot species were spotted on the center at an OD_600_ of 0.4. The plate diameter is 6 cm. Photos were taken after 48 h incubation at 30°C. **(C)** Pair-wise spot assays. 5 μl of the two species were plot next to each other at an OD_600_ of 1. Scale bar represents 3 mm. Photos were taken after 48 h incubation at 30°C. **(D)** Carbon source metabolic ability measured by phenotype microarrays. **Figure 5–figure supplement**. Growth curves in TSB medium.

We also assessed the carbon source metabolic ability by high-throughput phenotypic microarrays (Figure 5D). Ent and Pan were generalists that can utilize a wide variety of carbon sources. Other community members specialized in using amino acids and organic acids as carbon sources while having a limited ability to utilize nucleotides, sugar alcohols, and sugars. In the TSB medium, Ent and Chr had the highest growth rate and growth capacity than other isolates (Figure 5 – figure supplement), suggesting the high competitiveness of these two strains. Collectively, resource competition was widespread among SynCom members.

## Discussion

A key concern in SynCom research is to understand how microbial interactions affect community composition and productivity. In our study, we evaluated the contribution of each individual to community productivity and revealed the metabolic interactions. Our SynCom system identified Pan (*P. eucrina*) and *Chr* (*C. rhizoplanae*) as important drivers of community interaction networks through metabolic cross-feeding and resource competition. Subsequently, we predicted bacterial interactions by co-occurrence network analysis and tested the interactions in reduced communities.

Manipulating microbial consortia has a variety of applications; however, attempts to engineer them often fail to achieve the expected results owing to the unexpected effects of interactions within the community. To assemble a stable community, three main criteria should be considered: natural co-occurrence, tractability, and stability. Firstly, to represent a natural co-occurrence, the community should contain members that also co-exist in natural settings. Natural co-occurrence ensures a higher possibility of stable co-existence as well as higher natural relevance than artificial combination of isolates with unrelated origin. The model maize community (Niu et al., 2017), SXMP soil community (Ren et al., 2015), THOR community (Lozano et al., 2019), metal working fluids community (Piccardi et al., 2019), cheese rind community (Wolfe et al., 2014), and Yeast-LAB community (Ponomarova et al., 2017) were all constructed based on this criterium. Multiple species co-occurring in the natural microbiome could be pooled together to assemble an initial community. We assembled the initial community based on soil co-occurrence and co-existence in a laboratory biofilm. The second criterium is tractability. All the community members should be easily tractable by colony counting or qPCR, and the community should have an easily tractable trait, such as biofilm productivity. The third premise is a reduced community size. Species that cannot establish themselves in the initial community or species that share similar genetic backgrounds can be omitted from the community. The stability of a reduced community, the ability to maintain all the isolates, should be tested again at select time points of assembly. In this study, the community composition was tested by strain-specific qPCR. The reduced community was determined by network analysis and abundance: six correlated or abundant isolates were selected from the initial community.

We tested the positive or negative roles of the isolates as predicted by network analysis through the so-called removal strategy (where certain members were removed from the SynCom). Only 4 of the 6 isolates exhibited a predicted influence on community productivity. Therefore, although a co-occurrence network could serve as the first step for predicting bacterial interactions, this approach is insufficient for conclusively identifying keystone species and therefore requires experimental validation (Carlström et al., 2019; Faust, 2021). The removal strategy could be used to verify the role of the keystone species (Carlström et al., 2019; Niu et al., 2017).

After these three steps, further properties can be assessed according to the research interest, e.g., plant-growth-promoting properties, toxic fluid degrading ability, and stress resistance (Lee et al., 2014; Niu et al., 2017; Piccardi et al., 2019; Wang et al., 2021). Furthermore, potential interaction can be detected using spent medium growth assay, pair-wise growth experiments, and whole-genome metabolic modeling. Resource competition plays a major role in shaping bacterial communities, while metabolic exchanges promote group survival (Goldford et al., 2018; Zelezniak et al., 2015b). In the present study, we simulated the metabolic interactions using metabolic models and assessed their predictability using phenotypic assays. Both the metabolic model and the experiments suggested that the positive keystone species, Pan serves as a metabolic donor and supports the growth of other community members. On the contrary, the negative keystone species, Chr was the largest metabolic receiver in the community; it grew faster and accumulated higher growing capacity than other isolates except for Ent. Removal of this isolate decreased the metabolic resource overlap within the community, suggesting that it negatively affected community productivity through metabolic competition. These results indicate that metabolic interactions play a key role in determining the composition of communities, which is also observed in the maize model community (Krumbach et al., 2021). Importantly, five members of the SynCom were predicted to be donors by metabolic modeling, but only one species was experimentally confirmed in the applied experiments. This result could be attributed to the inherent uncertainty in genome-scale metabolic model reconstruction, such as the limited accuracy of genome annotation, the limitation of defined media, the lack of direct experimental measurements for most organisms, and the problems in network gap-filling (Bernstein et al., 2021). Thus, experimental data should be incorporated to curate the metabolic models.

One limitation of our study is the technical limitations to use identical media both for the biofilm experiments and during the metabolic assays. However, no biofilm is formed by these SynCom members in a minimal medium, while the metabolites exchange cannot be easily tested in a rich medium, like TSB. Nevertheless, the observed cross-feeding in the minimal medium is likely to occur in the biofilm, as reported in a dual-species biofilm community (Hansen et al., 2007). Furthermore, other mechanisms, such as nutrient availability, niche partitioning created by spatial and temporal heterogeneity, and trade-offs between nutrient acquisition and environmental tolerance, can also promote coexistence between species (Carlström et al., 2019; Estrela et al., 2021; Louca et al., 2018; Marchal et al., 2017). Various properties and mechanisms of bacterial interactions can be derived from the studies using model communities. For example, studies on the model Yeast-LAB community revealed that cross-feeding of amino acids might explain the co-existence of yeast and lactic acid bacteria in a variety of naturally fermented food and beverages (Ponomarova et al., 2017). Research on an SXMP soil community proposed the emergence of community intrinsic properties, such as enhanced biofilm formation and protection against grazing (Raghupathi et al., 2018; Ren et al., 2015). Further three-species biofilm community displayed enhanced resistance to antibiotics and SDS (Lee et al., 2014).

Although a potential practical application of the here developed SynCom is unclear, our research provides insights into how metabolic competition and cooperation simultaneously shape the community composition. The methodology of network co-occurrence analysis combined with qPCR quantification, metabolic modeling, and pair-wise interactions can be applied in diverse SynComs studies. Ultimately, such studies should translate to a deeper understanding of how microbial communities behave in their native environments, and this knowledge may be applied to wastewater treatment, disease suppression, and crop yield enhancement.

## Materials and methods

### Strains and genome sequencing

As shown in Table 1, eleven bacterial isolates derived from the cucumber rhizosphere (Sun et al., 2021) were selected for this study. These isolates were selected based on their co-existence in biofilms (see Results). Start inoculum was prepared by growing overnight culture in TSB medium, centrifuged (5,000g, 2 min), and resuspended in 0.9% NaCl solution to an optical density at 600 nm of one (OD_600_ ∼1). Multi-species start inoculum was prepared by mixing equal volumes of single-species initial inoculum.

The whole genomes of the selected isolates were sequenced by different companies at different times. The genomes of Aci, Com, Pxa, and Ste were sequenced using a combination of PacBio RS II and Illumina HiSeq 4000 sequencing platforms. The genome of Aci, Pxa, and Ste was sequenced at the Beijing Genomics Institute (BGI, Shenzhen, China). Four SMRT cells Zero-Mode Waveguide arrays of sequencing were used by the PacBio platform to generate the subreads set. PacBio subreads (length < 1 kb) were removed. The program Pbdagcon (https://github.com/PacificBiosciences/pbdagcon) was used for self-correction. Draft genomic unitigs, which are uncontested groups of fragments, were assembled using the Celera Assembler against a high-quality corrected circular consensus sequence subreads set. To improve the accuracy of the genome sequences, GATK (https://www.broadinstitute.org/gatk/) and SOAP tool packages (SOAP2, SOAPsnp, SOAPindel) were used to make single-base corrections. To trace the presence of any plasmid, the filtered Illumina reads were mapped using SOAP to the bacterial plasmid database (http://www.ebi.ac.uk/genomes/plasmid.html, last accessed July 8, 2016). Raw sequencing data and the assembled genome have been deposited to the National Center for Biotechnology Information (NCBI) under the BioProject accession number PRJNA593376, PRJNA762936, and PRJNA762715. The genome of Com was sequenced at Majorbio Bio-Pharm Technology Co., Ltd. Raw sequencing data and the assembled genome have been deposited to the NCBI under the BioProject accession number PRJNA762695.

The genomes of Bur, Chr, Ent, and Pan were sequenced using PacBio Sequel platform and Illumina NovaSeq PE150 at the Beijing Novogene Bioinformatics Technology Co., Ltd. Raw sequencing data and the assembled genomes have been deposited to the NCBI under the BioProject accession number PRJNA593683, PRJNA721858, PRJNA761942, and PRJNA762676. Pse was sequenced by (Sun et al., 2021). Genomes were automatically annotated by NCBI PGAP.

### Soil community co-existence prediction

To create a multi-species biofilm community, co-existence criterium was adopted. In parallel with our previous study (Sun et al., 2021), rhizosphere soil of cucumber soil was collected. Two types of soil were chosen: black soil and paddy soil. Bacterial abundance in the rhizosphere soil was roughly estimated by CFU counting on TSB agar plates. Equal volumes of soil suspension were mixed with *B. velezensis* SQR9 (OD_600_ ∼1) and inoculated in 2 ml of TSB liquid medium at a 1:100 ratio. Biofilms formed at the air-liquid interface and the solution underneath were collected separately after 24 h of incubation at 30°C. Each treatment had three biological replicates. The genomic DNA of the samples was extracted using an E.Z.N.A. Bacterial DNA Kit (Omega Bio-tek, Inc.) following the manufacturer’s instructions. Universal primers targeting the V3-V4 regions of the 16S rRNA gene were used to construct the DNA library for sequencing. Paired-end sequencing of bacterial amplicons was performed on the Illumina MiSeq instrument (300 bp paired-end reads). Raw sequencing data have been deposited to the NCBI SRA database under BioProject accession number PRJNA739098. Reads were processed using the UPARSE pipeline (http://drive5.com/usearch/manual/uparse_pipeline.html). The raw sequences were first trimmed at a length of 250 bp using the “fastx_truncate” command to discard shorter sequences. The paired-end reads were merged using the “fastq_mergepairs” command (less than 3 mismatches). High-quality sequences were then selected using the “fastq_filter” command (maximum error rate < 0.5%), and dereplicated using the “derep_fulllength” command. The singletons were removed using “unoise3” algorithm and chimeric sequences were removed using “uchime_ref” command with RDP database (RDP training set v16) (https://www.drive5.com/usearch/manual/sintax_downloads.html). The remaining sequences were clustered to operational taxonomic units (OTUs) based on 97% sequence similarity. Taxonomy assignment of the OTUs was classified using “sintax” algorithm (confidence threshold 0.6) with the RDP database. Finally, a rarefied OTU table was created using the USEARCH “otutab_norm” command at a depth of 10,000 reads per sample. The bacterial composition in the biofilm was visualized in Microsoft Office Excel 2019 to predict co-existence.

### Initial 11-species SynCom composition analysis

The eleven-species biofilm was cultivated by mixing 4 ml of start inoculum (1%) with 400 ml of tryptic soy broth (TSB) and incubating at 30°C for two to eight days. Biofilms formed at the air-liquid interface were collected on days 2, 4, 6, and 8. Each time point had eight biological replicates. The genomic DNA of the biofilm samples was extracted as previously described. 16S rRNA gene amplicon sequencing was conducted and analyzed as described above. The difference in the analysis is that the generated sequences were not clustered and were directly used to create the amplicon sequence variant (ASV) table. Taxonomy of the ASVs was assigned to the species with a reference database consisting of the full 16S rRNA gene sequences of the eleven species. The ASVs were rarified using the USEARCH “otutab_rare” command at a depth of 10,000 reads per sample. The ASV table was provided as supplementary table. Microbial co-occurrence networks were constructed to show the interactions among species during biofilm development. Spearman correlations among all taxa were calculated using the R *psych* package. Only edges with correlation scores > 0.6 were kept (*p* < 0.05, FDR-adjusted). Correlation networks were visualized via Gephi using the Fruchterman Reingold layout (Bastian et al., 2009).

### Reduced SynComs biomass quantification

The five- and six-species biofilms were grown in 6-well microtiter plates (VWR) insert with 100 μm sterile nylon mesh cell strainers (Biologix Cat #15-1100). 10 ml of TSB liquid medium and 100 μl of start inoculum were added. The plates were incubated for 24 h at 30 °C to allow the biofilm to grow on top of the nylon mesh cell strainer.

Biomass was defined by the fresh weight of biofilm. The cell strainer was taken out from the well, visible drops were removed with paper. Then the cell strainer was weighed. Pellicle fresh weight was the total weight minus the weight of the nylon mesh. Each treatment had six biological replicates.

### Reduced SynComs cell numbers quantification by qPCR

Strain-specific primers were designed for the selected six isolates. Genome comparison was performed using Roary (Page et al., 2015) to find out strain-specific single-copy genes of each isolate. Primers were designed to target these genes. qPCR was used to evaluate the specificity of the primers. Only primers that fulfill the following criteria were selected: (1) Selectively amplify target isolate but do not amplify noncognate isolates. (2) The CT values are low for targeting isolates, while high for non-targeting isolates. The CT values for non-targeting isolates are similar to that of water. (3) The melting curves display one peak for targeting isolates. The amplified fragments were ligated to PMD19T plasmids. Standard curves were generated using the plasmids containing corresponding fragments as templates. Detailed information on primers was provided in Supplementary information S1.

To quantify the cell numbers of each isolate within the reduced biofilms, 100 μl of the start inoculum was grown in six-well microtiter plates (VWR) with 10 ml TSB medium. A 100 μm sterile nylon mesh cell strainer and a Spectra Mesh Woven Filter (Fisher Scientific, Spectrum 146488) were put inside. The mesh was manually cut into 1.5 cm^2^ squares and autoclaved. The mesh ensured equal sampling of biofilm. After 24 or 36 h of pellicle development, the nylon mesh cell strainer was taken out, the inner filter was transferred to a 1.5 ml microcentrifuge tube, stored at −80 °C for following DNA extraction. The genomic DNA of the biofilm samples was extracted as previously described. qPCR was performed with Applied Biosystems Real-Time PCR Instrument. Reaction components are as follows 7.2 μl H_2_O, 10 μl 2× ChamQ SYBR qPCR Master Mix (Vazyme), 0.4 μl 10 μM of each primer and 2 μl template DNA. The PCR programs were carried out under the following conditions: 95 °C for 10 min, 40 cycles of 95 °C for 30 s, 60 °C for 45 s, followed by a standard melting curve segment. Each treatment had six biological replicates, and each sample was run in triplicates (technical replicates).

### Pair-wise assay

The direct competition of these isolates against each other was evaluated using the spot-on-lawn assay and the pair-wise spot assay. Spot-on-lawn assay: 5 ml of lawn species (OD_600_ ∼ 0.02) grown in TSB medium was spread onto a 25 ml TSB plate (1.5% agar) and removed by pipetting. Plates were dried for 20 min. 5 μl of spot species (OD_600_∼ 0.4) grown in TSB medium was spotted on the center of the plates. Pair-wise spot assay: 5 μl of the dual-species (OD_600_∼ 1) grown in TSB medium were spotted on the TSB plate (1.5% agar) at 5 mm between the center of each colony. Plates were grown at 30 °C and imaged at 48 h. The experiments were performed twice, each experiment had three replicate plates.

### Growth curve assay

Growth in the TSB medium was evaluated by growth curve assay. 2 μl of isolates (OD_600_ ∼1) was inoculated to 200 μl TSB medium in a 10×10 well Honeycomb Microplate. OD_600_ was measured every 30 minutes at 30 °C with Bioscreen C Automated Microbiology Growth Curve Analysis System. The growth was measured for 24 hours. Each treatment has five replicates.

The potential growth promotion of the bacterial metabolites to another species was evaluated using a spent medium growth curve assay. Donor bacteria were grown in the M9 medium with 0.2% glucose till the glucose was under detection. The consumption of glucose was measured using the Glucose GO Assay Kit (Sigma). The cell culture was spun down, then the spent medium was filter-sterilized and directly used as the medium of growth curve assay. 2 μl of recipient species (OD_600_∼ 1) was inoculated to 200 μl spent medium or M9 glucose medium in a 10×10 well Honeycomb Microplate. OD_600_ was measured every 30 minutes at 30 °C for 48 hours. Each treatment has five replicates. The carrying capacity (maximum population size) was compared.

### Genome-scale metabolic modeling

Metabolic model of the six-species reduced SynCom was reconstructed using the CarveMe pipeline (Machado et al., 2018). The M9 glucose medium was used as input medium for reconstructing genome-scale metabolic models. The quality of the metabolic models was validated using MEMOTE (Lieven et al., 2020). The metabolic interaction potential and metabolic resource overlap for each community were analyzed using SMETANA (Zelezniak et al., 2015; Zorrilla et al., 2021). The simulated cross-feeding results were summarized as SMETANA score, which estimates the strength of metabolic exchanging (Zelezniak et al., 2015).

### Carbon source metabolic activity measurement

PM1 (BIOLOG Cat #13101) and PM2 (BIOLOG Cat #13102) phenotypic microarrays were used to assess the carbon source utilization ability of the community members (Bochner et al., 2001). The assays were performed following the manufacturer’s instructions. Briefly, 100 μl of diluted cell suspension of each species mixed with the BiOLOG redox dyes were added to each well of the PM plates. Water mixed with dyes were used as negative controls. All of the plates were then incubated at 30 °C for up to 48 h. If the species could utilize the carbon source in a well, the colorless tetrazolium dye will be reduced to purple formazan by cell respiration. The color changes were measured by an endpoint absorbance at 590 nm with a microplate reader. The variable level of color changes indicates the carbon source metabolic activity.

### Data analysis and figures

All the data needed to evaluate the conclusions in this paper are provided in the figures. Source data related to this paper would be available online upon publication. Data were analyzed using R v4.1.3 (R Core Team, 2022) in the RStudio v2022.02.0+443. Statistical analysis methods were described in figure legends. Plots were generated using Microsoft Office Excel 2019 (stacked bar plot), R *ggplot2* (Wickham, 2016), *ggpubr* (Kassambara, 2020), *ggalluvial* (Brunson JC, 2020), *pheatmap* (Kolde, 2019) packages, and Adobe Illustrator CC 2020 (Adobe Inc.). Schematic diagrams were generated using BioRender (https://biorender.com/).

## Acknowledgments

This work was financially supported by the National Nature Science Foundation of China (31972506, 31972512 and 42090064), the National Key Research and Development Program (2022YFF1001800, 2021YFD1900300 and 2022YFD1500041). XS was supported by a Chinese Scholarship Council fellowship. ÁTK was supported by the Danish National Research Foundation (DNRF137) for the Center for Microbial Secondary Metabolites and the Novo Nordisk Foundation via the INTERACT project (grant number NNF19SA0059360). During her unexpected trap in Denmark caused by the pandemic of Omicron, author XS is extremely grateful to ZX for his financial support, and to Danish guitarist Niklas Johansen for his weekly lessons and constant spiritual support.

## Author contributions

XS, JX, QH, ZX, ÁTK designed the study, XS, JX, RX performed the experiments. DZ, WX, WW performed the metabolic modeling. XS and JX analyzed the data and created the figures. XS wrote the first draft of the manuscript, ZX, ÁTK, RZ, QS revised the manuscript.

---

**Figure 1–figure supplement 1.**
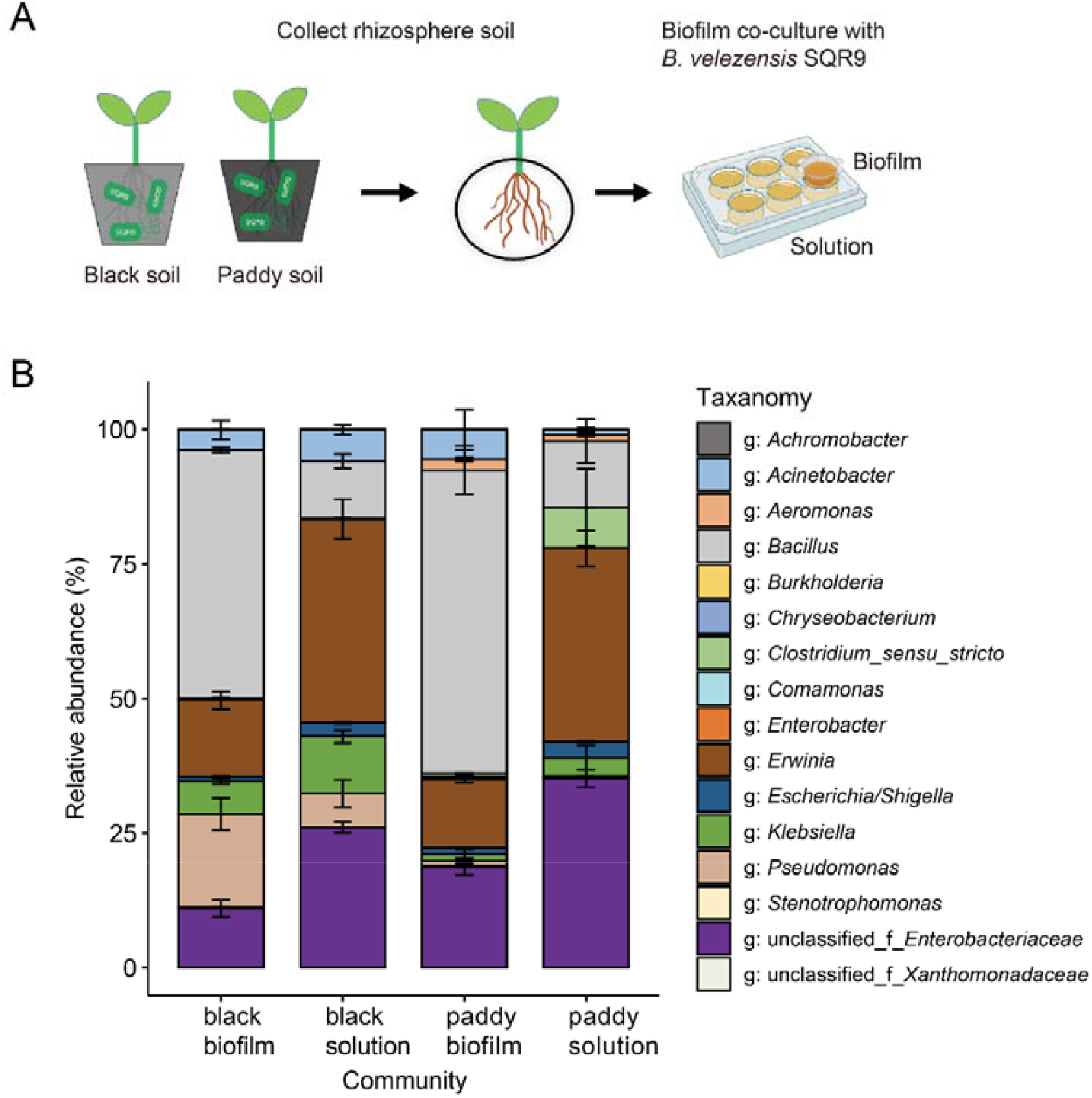
Origin of the initial 11 isolates. **(A)** Schematic diagram of the experimental setup. Rhizospheric black soil and paddy soil were collected from cucumber plants. The soil microbiota were co-cultivated with *B. velezensis* SQR9 at 30°C in TSB medium to form pellicle biofilms. After 24 hours of cultivation, the pellicle and the solution underneath were collected separately. The samples were sent for 16S rDNA amplicon sequencing. **(B)** Microbiome composition of the biofilm and solution. Data presented are the mean ±sd. n =3. Based on OTU clustering and the RDP database taxonomic classification, 15 genera and 2 families were identified. Eleven matching isolates were selected from our laboratory bacteria collection (Sun et al., 2021).

**Figure 1–figure supplement 2.**
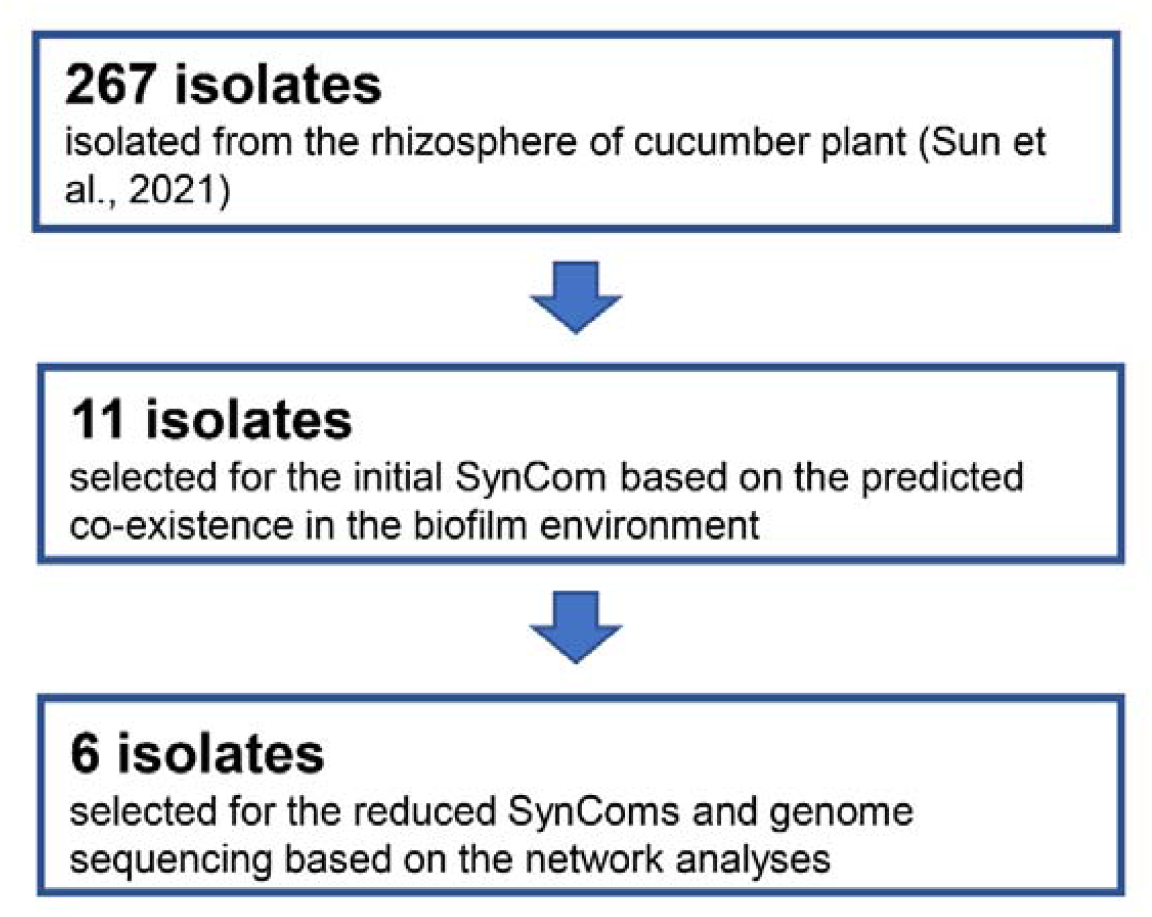
A flow chart of isolates used for the study, their origin and selection criteria.

**Figure 2–figure supplement.**
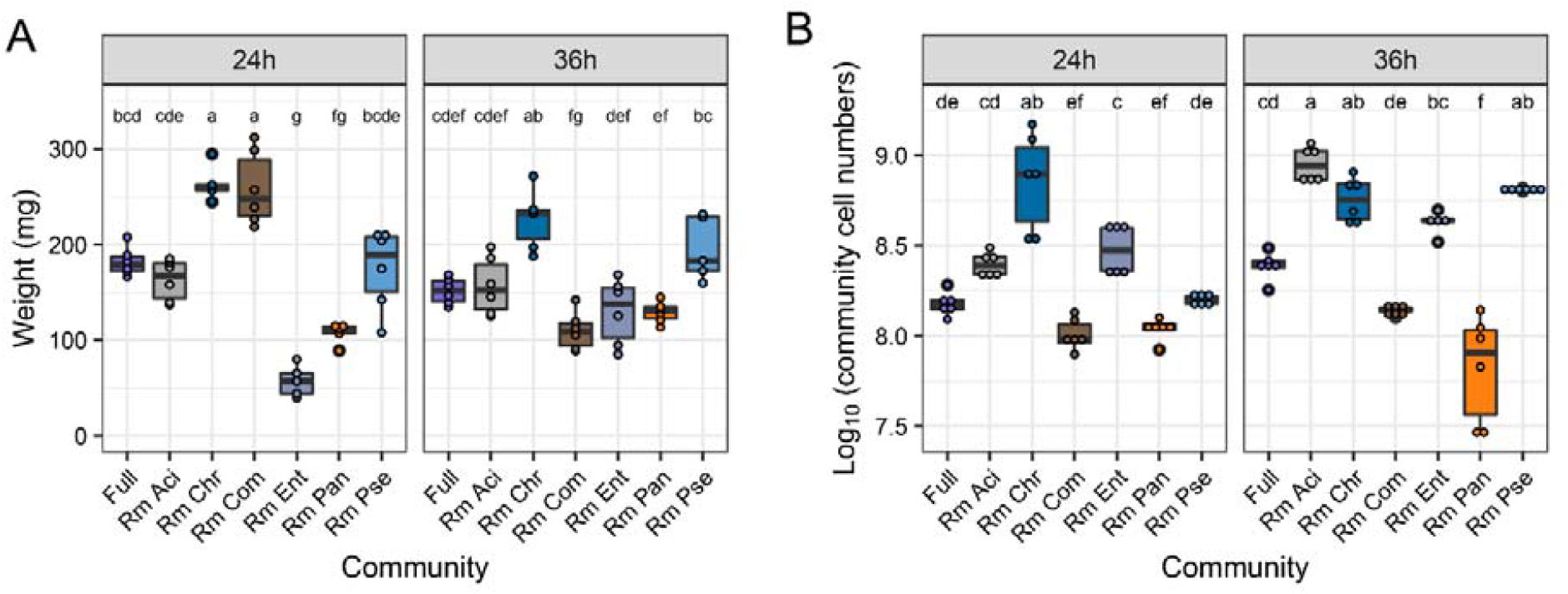
Productivity of the reduced SynComs. **(A)** Biofilm weight. **(B)** Cell numbers. Different letters indicate significant difference by one-way ANOVA, Tukey’s test. Data presented are the mean ±sd. n =6.

**Figure 3–figure supplement.**
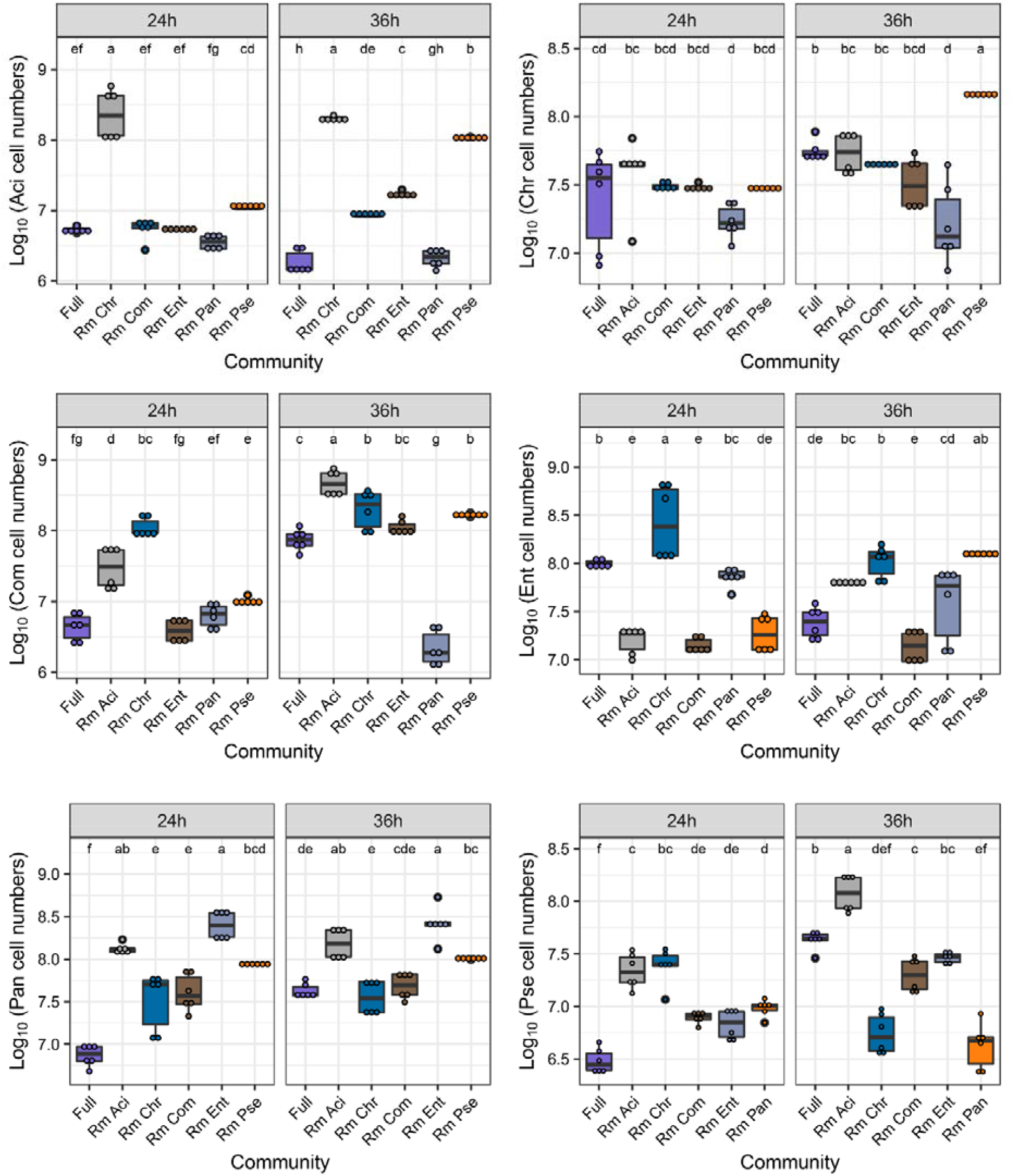
Individual cell numbers in different SynComs. Different letters indicate significant difference by one-way ANOVA, Tukey’s test. Data presented are the mean ±sd. n =6.

**Figure 4–figure supplement.**
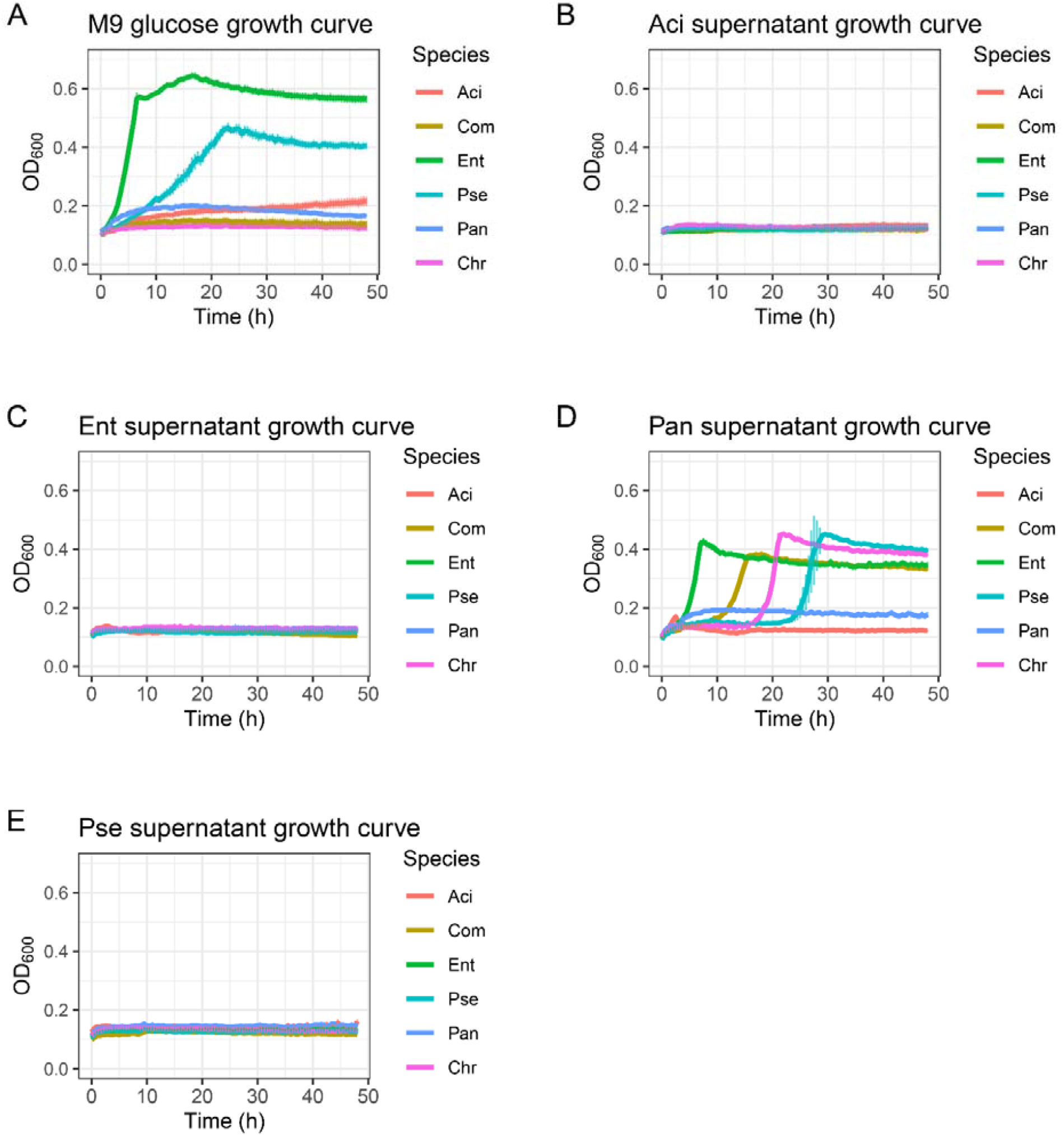
Growth curves. (A) Growth curves in M9 medium with 0.2% glucose as carbon source. Water was set as control inoculation. (B) Growth curves in Aci spent medium. (C) Growth curves in Ent spent medium. (D) Growth curves in Pan spent medium. (E) Growth curves in Pse spent medium. Data presented are the mean ±sd. n =5.

**Figure 5–figure supplement.**
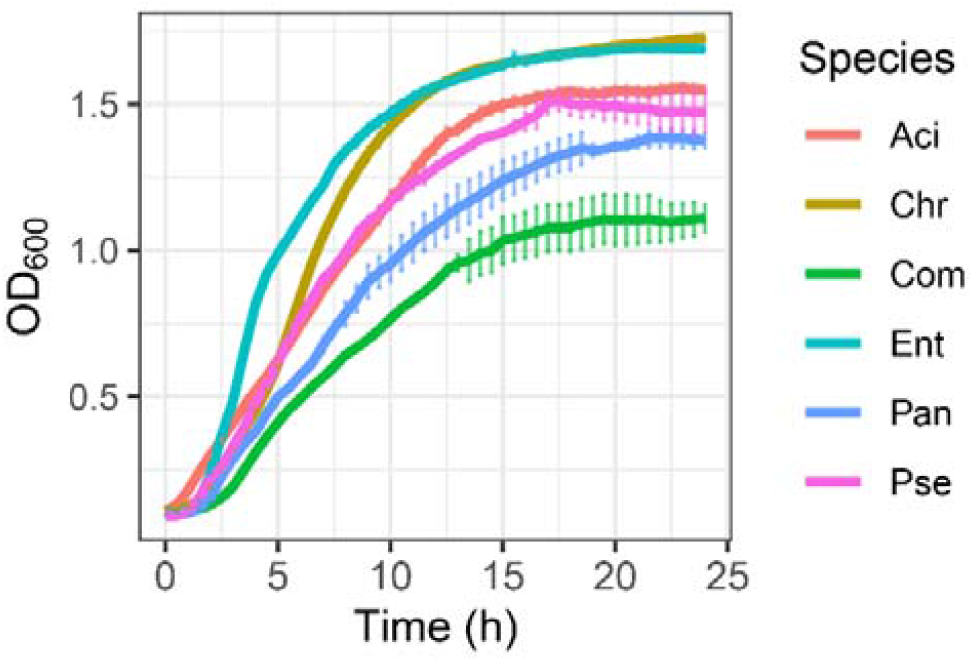
Growth curves in TSB medium. Data presented are the mean ±sd. n =5.

