## Supplementary information for "Keystone species determine the productivity of synthetic microbial biofilm communities"

### Slide 1
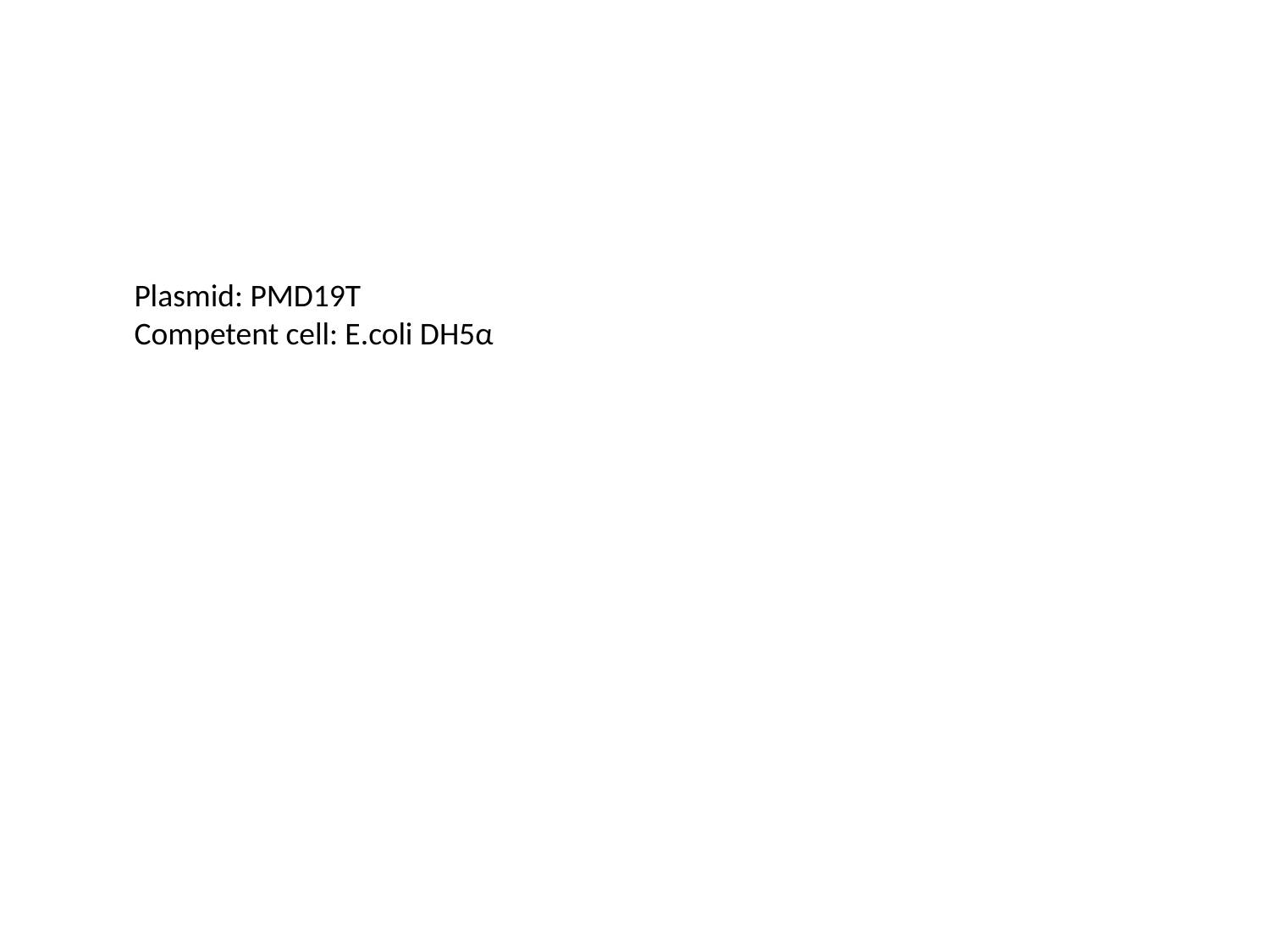

Plasmid: PMD19T
Competent cell: E.coli DH5α

### Slide 2
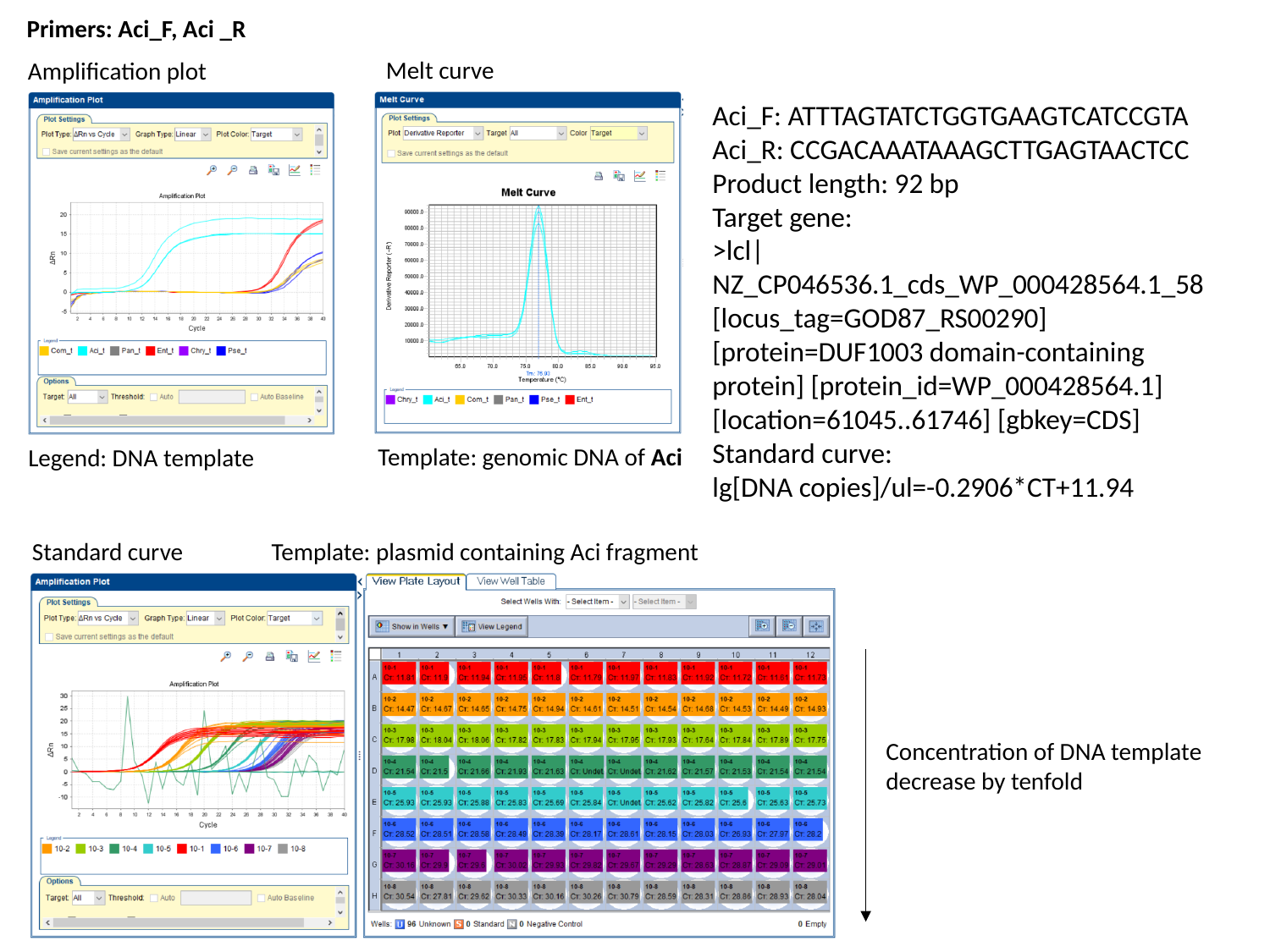

Primers: Aci_F, Aci _R
Melt curve
Amplification plot
Aci_F: ATTTAGTATCTGGTGAAGTCATCCGTA
Aci_R: CCGACAAATAAAGCTTGAGTAACTCC
Product length: 92 bp
Target gene:
>lcl|NZ_CP046536.1_cds_WP_000428564.1_58 [locus_tag=GOD87_RS00290] [protein=DUF1003 domain-containing protein] [protein_id=WP_000428564.1] [location=61045..61746] [gbkey=CDS]
Standard curve:
lg[DNA copies]/ul=-0.2906*CT+11.94
Template: genomic DNA of Aci
Legend: DNA template
Standard curve
Template: plasmid containing Aci fragment
Concentration of DNA template decrease by tenfold

### Slide 3
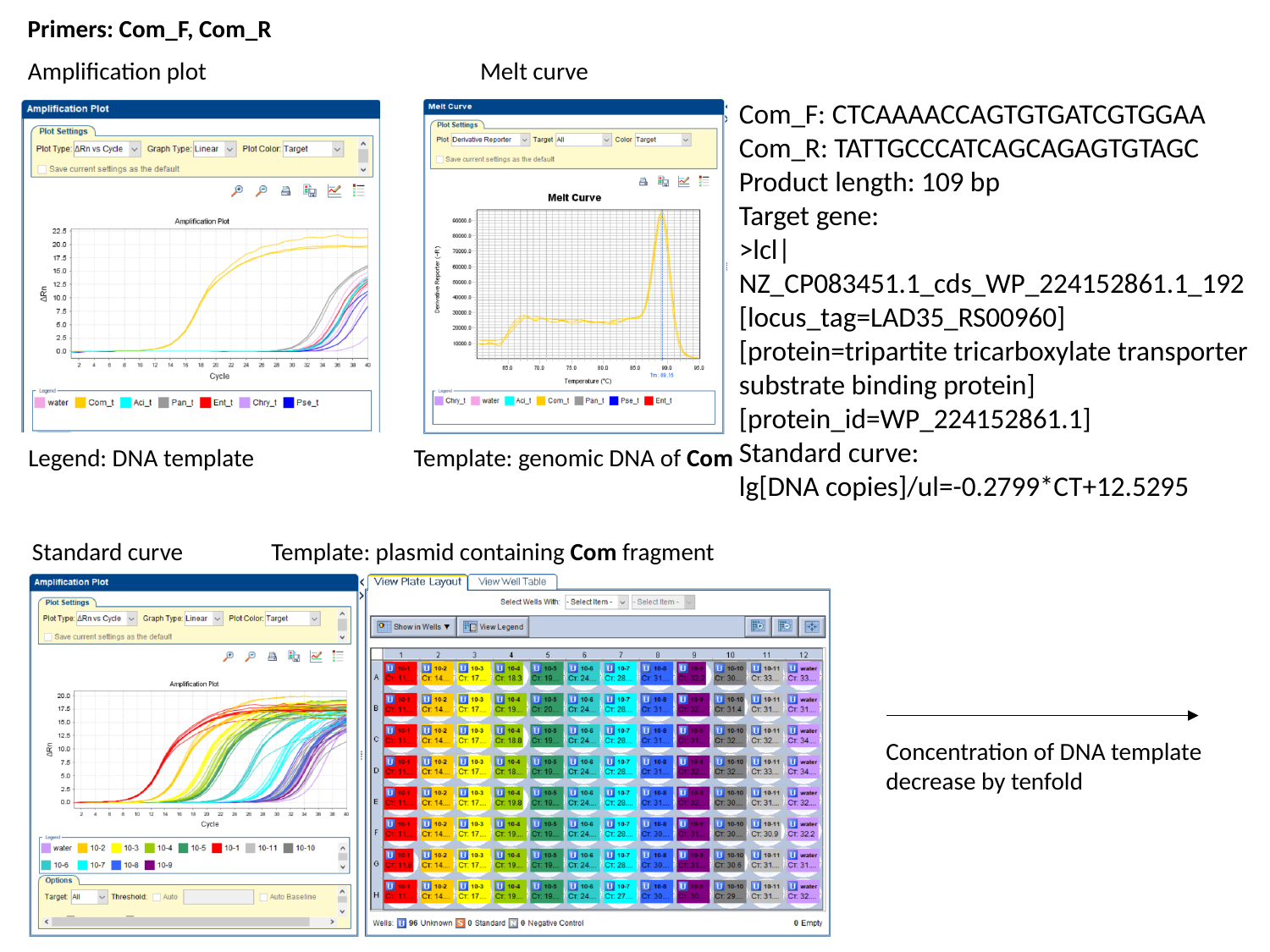

Primers: Com_F, Com_R
Amplification plot
Melt curve
Com_F: CTCAAAACCAGTGTGATCGTGGAA
Com_R: TATTGCCCATCAGCAGAGTGTAGC
Product length: 109 bp
Target gene:
>lcl|NZ_CP083451.1_cds_WP_224152861.1_192 [locus_tag=LAD35_RS00960] [protein=tripartite tricarboxylate transporter substrate binding protein] [protein_id=WP_224152861.1]
Standard curve:
lg[DNA copies]/ul=-0.2799*CT+12.5295
Legend: DNA template
Template: genomic DNA of Com
Standard curve
Template: plasmid containing Com fragment
Concentration of DNA template decrease by tenfold

### Slide 4
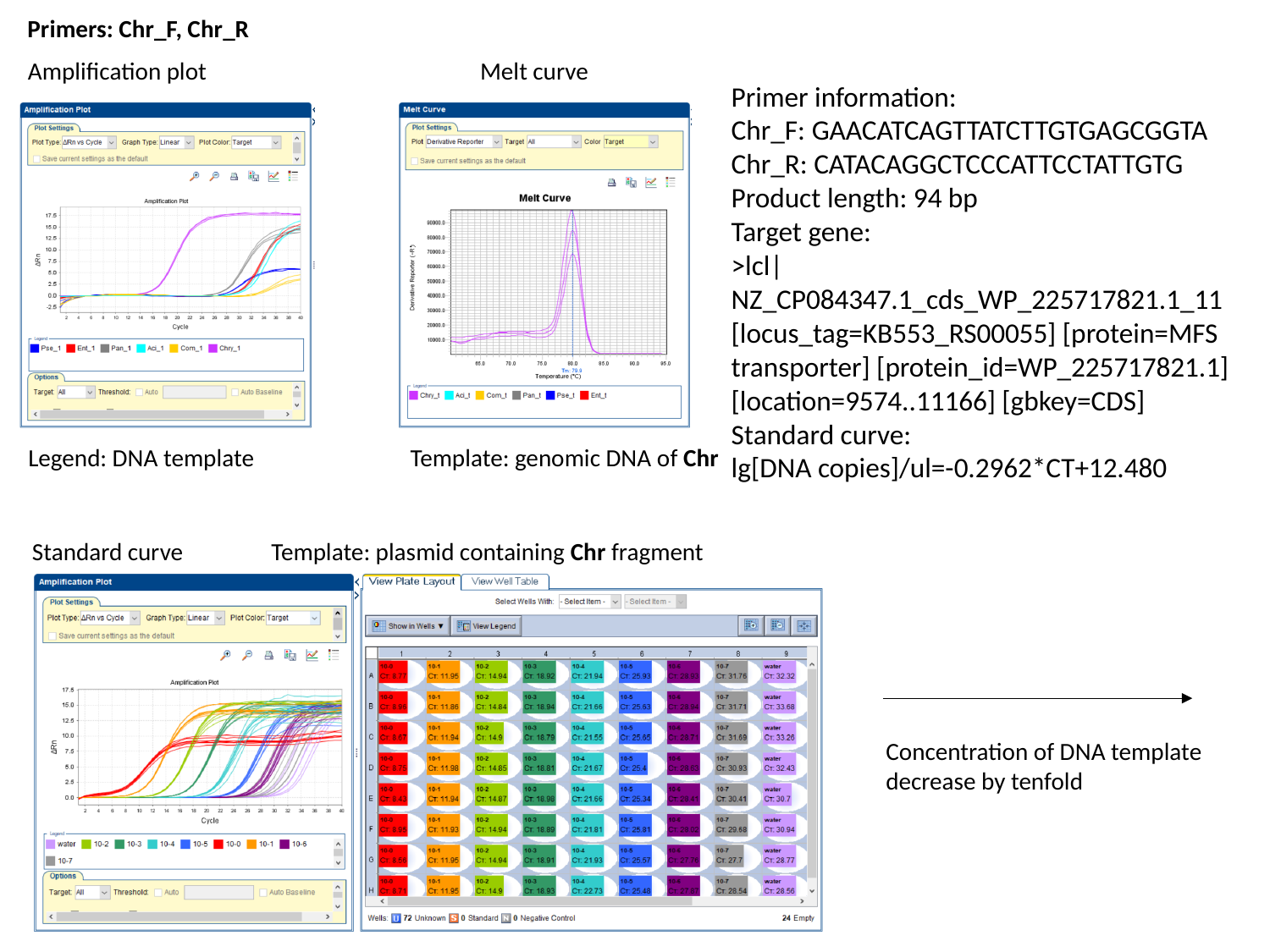

Primers: Chr_F, Chr_R
Amplification plot
Melt curve
Primer information:
Chr_F: GAACATCAGTTATCTTGTGAGCGGTA
Chr_R: CATACAGGCTCCCATTCCTATTGTG
Product length: 94 bp
Target gene:
>lcl|NZ_CP084347.1_cds_WP_225717821.1_11 [locus_tag=KB553_RS00055] [protein=MFS transporter] [protein_id=WP_225717821.1] [location=9574..11166] [gbkey=CDS]
Standard curve:
lg[DNA copies]/ul=-0.2962*CT+12.480
Legend: DNA template
Template: genomic DNA of Chr
Standard curve
Template: plasmid containing Chr fragment
Concentration of DNA template decrease by tenfold

### Slide 5
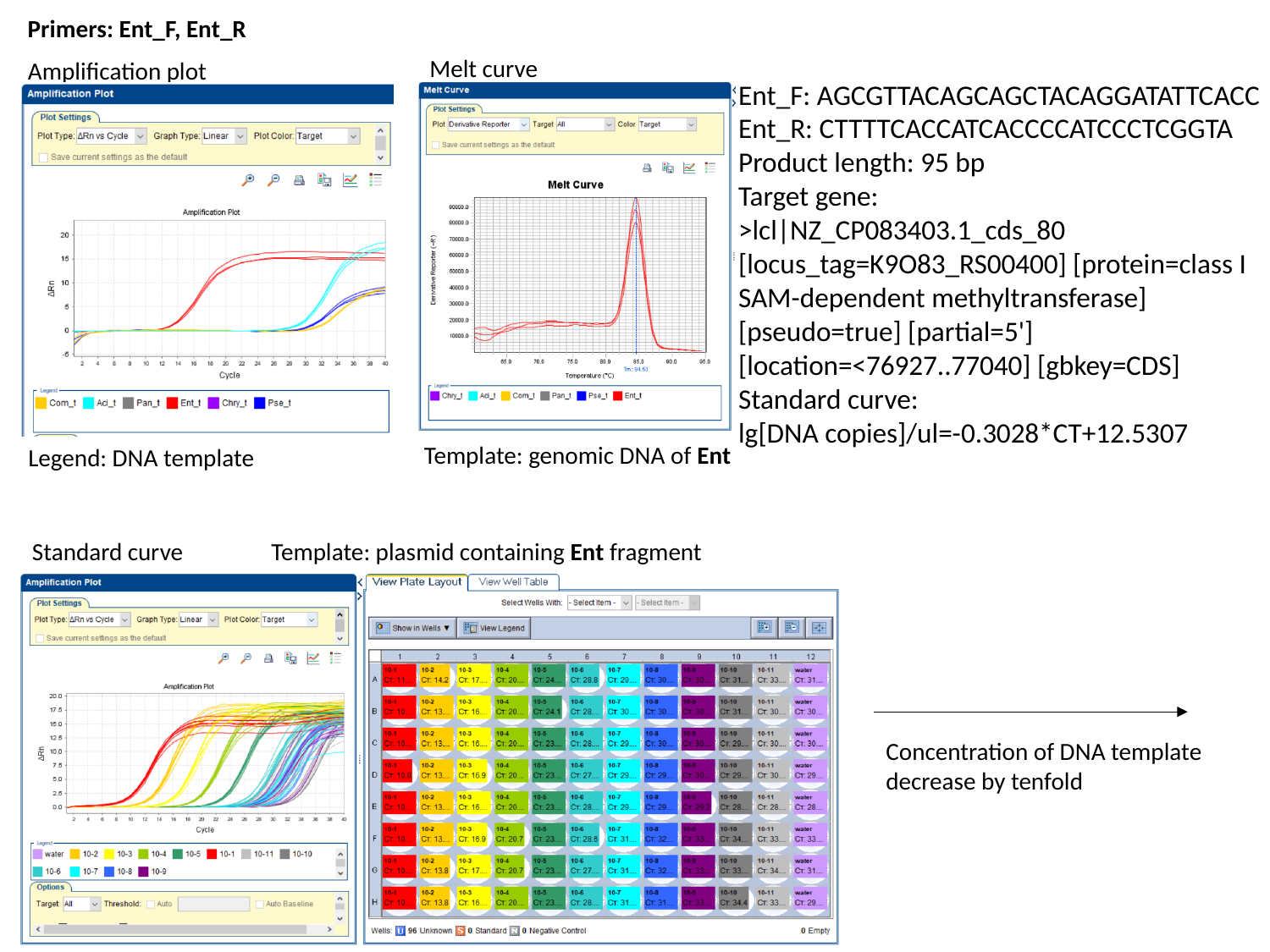

Primers: Ent_F, Ent_R
Melt curve
Amplification plot
Ent_F: AGCGTTACAGCAGCTACAGGATATTCACC
Ent_R: CTTTTCACCATCACCCCATCCCTCGGTA
Product length: 95 bp
Target gene:
>lcl|NZ_CP083403.1_cds_80 [locus_tag=K9O83_RS00400] [protein=class I SAM-dependent methyltransferase] [pseudo=true] [partial=5'] [location=<76927..77040] [gbkey=CDS]
Standard curve:
lg[DNA copies]/ul=-0.3028*CT+12.5307
Template: genomic DNA of Ent
Legend: DNA template
Standard curve
Template: plasmid containing Ent fragment
Concentration of DNA template decrease by tenfold

### Slide 6
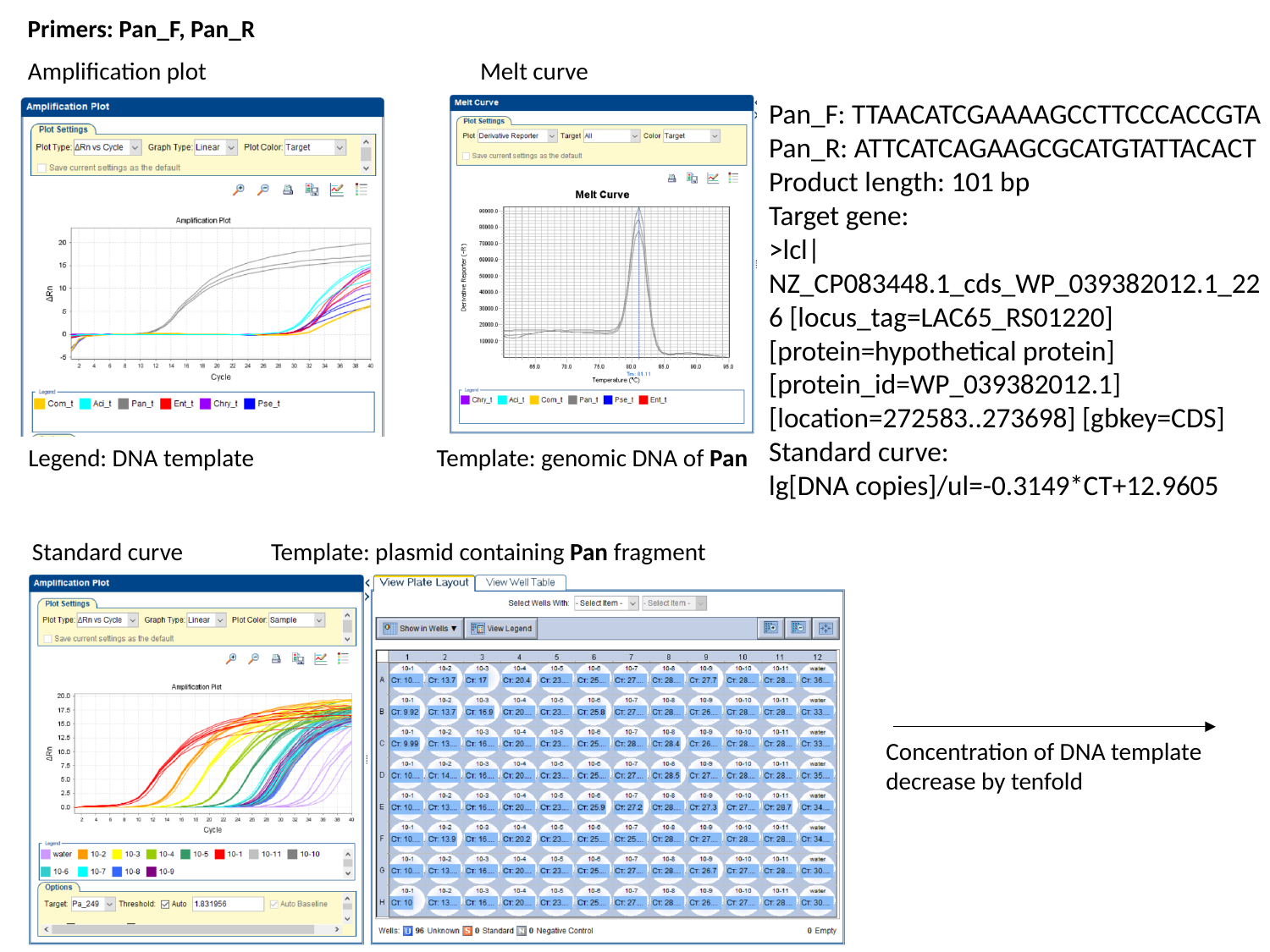

Primers: Pan_F, Pan_R
Amplification plot
Melt curve
Pan_F: TTAACATCGAAAAGCCTTCCCACCGTA
Pan_R: ATTCATCAGAAGCGCATGTATTACACT
Product length: 101 bp
Target gene:
>lcl|NZ_CP083448.1_cds_WP_039382012.1_226 [locus_tag=LAC65_RS01220] [protein=hypothetical protein] [protein_id=WP_039382012.1] [location=272583..273698] [gbkey=CDS]
Standard curve:
lg[DNA copies]/ul=-0.3149*CT+12.9605
Template: genomic DNA of Pan
Legend: DNA template
Standard curve
Template: plasmid containing Pan fragment
Concentration of DNA template decrease by tenfold

### Slide 7
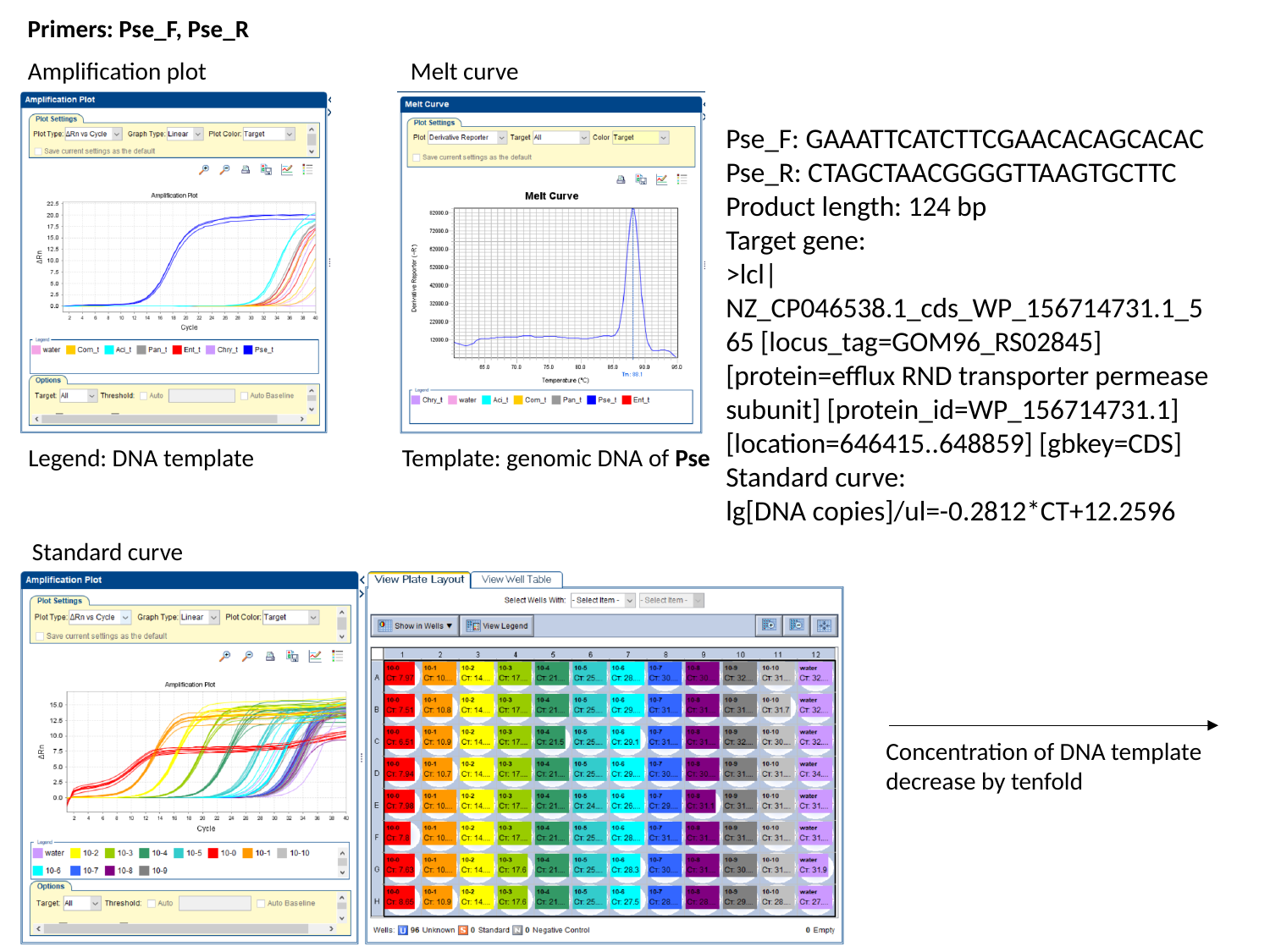

Primers: Pse_F, Pse_R
Amplification plot
Melt curve
Pse_F: GAAATTCATCTTCGAACACAGCACAC
Pse_R: CTAGCTAACGGGGTTAAGTGCTTC
Product length: 124 bp
Target gene:
>lcl|NZ_CP046538.1_cds_WP_156714731.1_565 [locus_tag=GOM96_RS02845] [protein=efflux RND transporter permease subunit] [protein_id=WP_156714731.1] [location=646415..648859] [gbkey=CDS]
Standard curve:
lg[DNA copies]/ul=-0.2812*CT+12.2596
Legend: DNA template
Template: genomic DNA of Pse
Standard curve
Concentration of DNA template decrease by tenfold
